# Medroxyprogesterone acetate mediated alteration in the vaginal microbiota and microenvironment in a Kenyan sex worker cohort

**DOI:** 10.1101/483180

**Authors:** Jocelyn M. Wessels, Julie Lajoie, Maeve I. J. Hay Cooper, Kenneth Omollo, Allison M. Felker, Danielle Vitali, Haley A. Dupont, Philip V. Nguyen, Kristen Mueller, Fatemeh Vahedi, Joshua Kimani, Julius Oyugi, Juliana Cheruiyot, John N. Mungai, Alexandre Deshiere, Michel J. Tremblay, Tony Mazzulli, Jennifer C. Stearns, Ali A. Ashkar, Keith R. Fowke, Michael G. Surette, Charu Kaushic

## Abstract

The hormonal contraceptive Medroxyprogesterone Acetate (MPA) is associated with increased risk of Human Immunodeficiency Virus (HIV), via incompletely understood mechanisms. Increased diversity in the vaginal microbiota modulates genital inflammation and is associated with increased HIV-1 acquisition. However, the effect of MPA on diversity of the vaginal microbiota is relatively unknown. In a cohort of female Kenyan sex workers, negative for sexually transmitted infections (STIs), with Nugent Scores <7 (N=58 of 370 screened), MPA correlated with significantly increased diversity of the vaginal microbiota as assessed by 16S rRNA gene sequencing. MPA was also significantly associated with low vaginal glycogen and α-amylase, factors implicated in vaginal colonization by lactobacilli, bacteria believed to protect against STIs. Furthermore, increased diversity of the vaginal microbiota correlated with activation of vaginal HIV-1 target cells. Results were recapitulated in humanized mice where MPA treatment was associated with increased diversity of the vaginal microbiota, low glycogen, and enhanced HIV-1 susceptibility. Together these results suggest MPA-induced hypo-estrogenism may alter key metabolic components necessary for vaginal colonization by certain bacterial species including lactobacilli, and allow for greater bacterial diversity in the vaginal microbiota. Bacterial diversity in the vaginal microbiota correlates with activation of HIV-1 target cells, which might thus contribute to enhanced susceptibility to HIV-1.

## Introduction

Meta-analyses suggest the injectable progestin-based contraceptive Depot-Medroxyprogesterone Acetate (DMPA) increases heterosexual acquisition of Human Immunodeficiency Virus (HIV-1) 1.4-fold (1), while a prospective study reported women using injectable progestins were at 3.5 times increased risk of HIV acquisition compared to women not using long-term hormonal contraceptives (2). These statistics are particularly troubling because DMPA is a popular contraceptive in Africa (3), where HIV-1 prevalence is greatest. Studies have suggested several biological mechanisms, including hypo-estrogenism, by which DMPA might enhance susceptibility to HIV-1 and other sexually transmitted infections (STIs) (4). However, few (5-10) examine the relationship between hormonal contraceptives and the vaginal microbiota (VMB), even though diverse VMB, low in *Lactobacillus* species, confers a 4-fold increased risk of HIV-1 (11). Herein, we determined the effect of MPA (active component of DMPA) on several factors within the vaginal microenvironment of Kenyan sex workers, including diversity of the VMB.

The VMB is a bacterial community lining vaginal epithelial cells (12). Unlike the diverse gut microbiota, the VMB is generally low in diversity, and five community state types, including four dominated by a *Lactobacillus* species have been described (12). Two factors thought to maintain vaginal lactobacilli are glycogen, stored in epithelial cells and available as free glycogen in vaginal fluid (13-15), and α-amylase, an enzyme that catabolizes glycogen for use by lactobacilli and other bacteria (16) as energy. Similar to the gut, the VMB modifies immunity. The VMB alters inflammation in the female genital tract (17), and is implicated in reproductive health and disease. Several factors including ethnicity (12), STIs (18), bacterial vaginosis (BV) (19, 20), and sex work (21) have been reported to impact VMB composition. This study was designed to examine the effect of hormonal contraceptives on the VMB of healthy, asymptomatic Kenyan sex workers to determine if DMPA is associated with changes in the VMB and vaginal microenvironment that might impact their susceptibility to HIV-1. Importantly, we chose to exclude women with BV because BV is a clinical condition reported to profoundly overshadow the effect of hormones on vaginal biomarkers of microbial health (glycosidases, lectins) in cervico-vaginal lavage (CVL) (22, 23). Furthermore, we (21) and others (19, 20) have demonstrated by 16S ribosomal RNA (rRNA) gene sequencing that women diagnosed with BV by Nugent scoring have inherently diverse VMBs, but women with high diversity VMBs do not always have Nugent scores indicative of BV. Therefore, because symptomatic BV is a clinical condition that could potentially confound the relationship between DMPA and diversity of the VMB, and alter vaginal biomarkers of microbial health the present study excluded women with Nugent scores 7-10, which is used to diagnose clinical BV.

Meta-analyses find DMPA correlates with increased HIV-1 susceptibility in women (1), and that women with diverse VMB have increased susceptibility to HIV-1 (11). Based on these observations, we hypothesized that MPA will increase diversity of the VMB, which in turn will affect HIV-1 risk. Herein we examine the effect of MPA on the VMB, vaginal glycogen, and vaginal α-amylase in healthy, asymptomatic Kenyan sex workers. We also explore the relationship between bacterial diversity, vaginal cytokines, and HIV-1 target cells. Our overall aim was to perform bacterial 16S rRNA gene sequencing of the VMB of sex workers using DMPA, oral contraceptives (OCPs), or not using hormonal contraceptives (NH; proliferative phase of the menstrual cycle), and assess the effect of hormonal contraceptives on the vaginal microenvironment and diversity of the VMB, as they relate to HIV-1 susceptibility. We found DMPA associated with increased VMB diversity, diversity correlated with changes in HIV-1 target cells, and that vaginal glycogen and α-amylase were lower in DMPA-treated sex workers. We experimentally recapitulated the results of our clinical study in humanized mice, and demonstrated that estrogen suppressed VMB diversity, and DMPA enhanced HIV-1 susceptibility. Results suggest MPA-induced hypo-estrogenism alters key metabolic products (ie. glycogen, and α-amylase) important for vaginal colonization by protective bacterial species like lactobacilli, and the change in substrates might allow for greater bacterial diversity. Diversity of the VMB positively correlates with activation of HIV-1 target cells, indicating the enhanced bacterial diversity in sex workers on DMPA might contribute to increased susceptibility to HIV-1.

## Results

### DMPA is associated with vaginal microbial diversity in healthy, asymptomatic Kenyan sex workers

Our primary objective was to examine the effect of DMPA on VMB diversity. We enrolled a select group of Kenyan sex workers, according to strict inclusion/exclusion criteria listed in Methods. As several factors including STIs and BV impact diversity of VMB, women with STIs and/or BV were excluded. Fifty-eight (N=58) women met study criteria (Supplemental Figure 1). Circulating MPA, estradiol, and progesterone were quantified in study participants (Supplemental Figure 2A, B, C). As previously reported, use of DMPA was associated with hypo-estrogenism v(4, 24, 25); sex workers on DMPA had significantly lower levels of circulating estradiol compared to NH women (Supplemental Figure 2B). We also found DMPA was associated with significantly lower levels of circulating progesterone, as compared to NH sex workers and those on OCPs (Supplemental Figure 2C).

We sought to determine the effect of hormonal contraceptives on diversity of the VMB. Alpha-diversity was assessed in sex workers not on hormonal contraceptives (NH, N=22, proliferative phase of the menstrual cycle), on OCPs (N=14), and on DMPA (N=22), by observed species (richness), Chao1 (richness), and Shannon Diversity Index (evenness and richness). While significant differences in observed species (Figure 1A) were initially observed during Kruskal-Wallis one-way ANOVA, no significant differences were identified during post-hoc testing. While Chao1 richness (Figure 1B), was significantly higher in the VMB of sex workers on DMPA versus those on OCPs, no significant differences were observed between NH women and women on DMPA. However, the VMB of sex workers on DMPA was significantly more diverse than NH (Figure 1C,1D), and OCP women (Figure 1C), as seen in Shannon Diversity rarefaction curves. To rule out that the bacterial diversity associated with DMPA might be due to a disproportionate number of women with intermediate (4-6) Nugent Scores, this group was removed from the analysis and Shannon Diversity was plotted in women with Nugent Scores 0-3 (Supplemental Figure 3; NH, N=18; OCP, N=13; DMPA, N=17). Even in this subset, the Shannon Diversity Index was significantly greater in sex workers on DMPA than NH, suggesting that DMPA is associated with enhanced bacterial diversity in the VMB of healthy, asymptomatic Kenyan sex workers.

**Figure 1:**
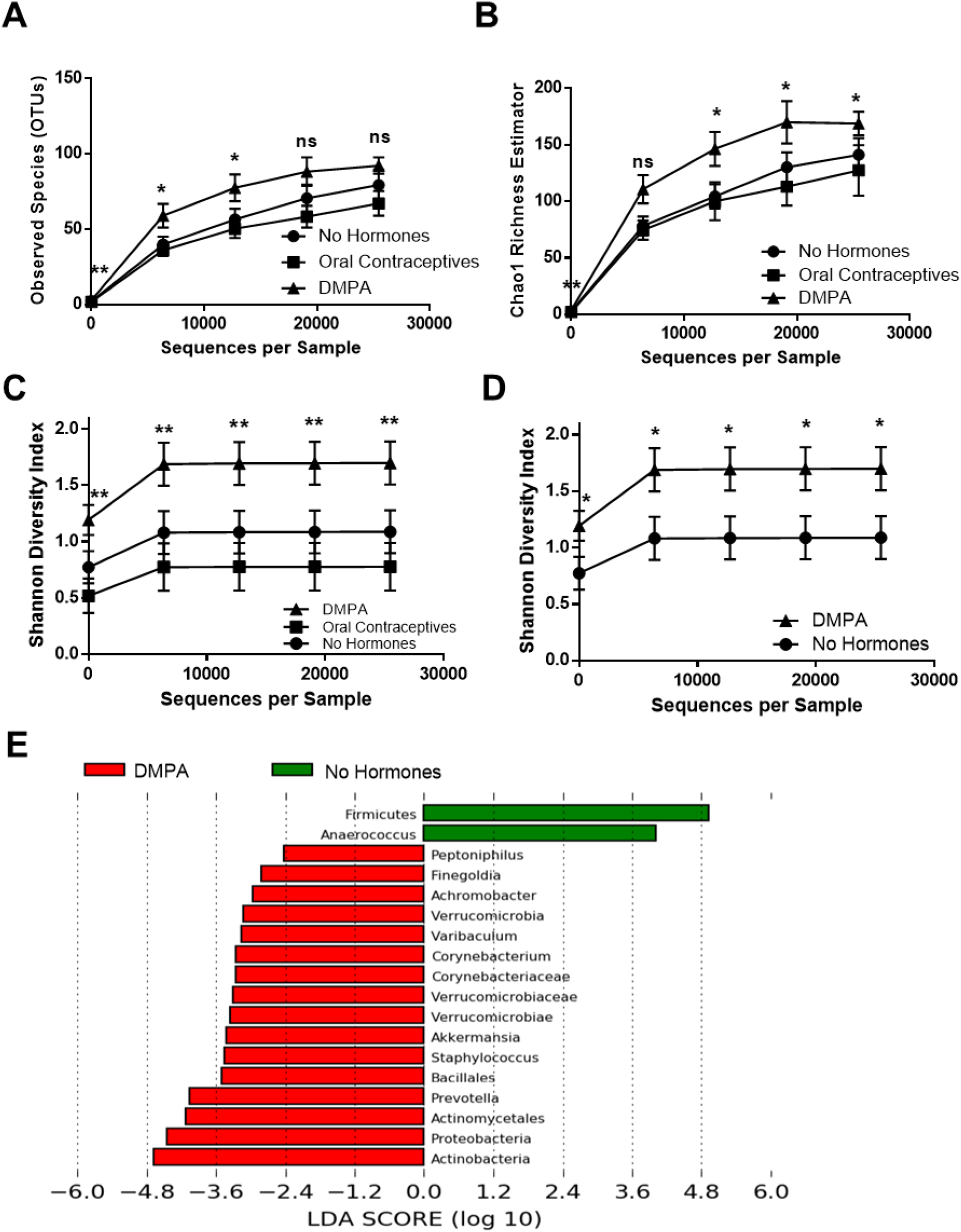
DMPA Associated with Vaginal Bacterial Diversity in Kenyan Sex Workers. Three alpha-diversity metrics were used to compare bacterial richness and evenness within the vaginal microbiota of Kenyan sex workers not on hormonal contraceptives (proliferative phase of menstrual cycle, N=22), on oral contraceptives (N=14), and on DMPA (N=22), from the same geographical region. As BV is a clinical condition that might potentially confound the relationship between DMPA and bacterial diversity, the present study excluded sex workers with Nugent scores 7-10. (**A**) Significant differences in Observed Species were initially observed during Kruskal-Wallis one-way ANOVA, however no significant differences were subsequently identified during post-hoc testing. (**B**) Chao1 richness was significantly higher in the VMB of sex workers on DMPA versus those on OCPs (P≤0.05; Kruskal-Wallis one-way ANOVA), yet no significant differences were observed between NH sex workers and those on DMPA. (**C**) Sex workers on DMPA had the greatest bacterial diversity at all levels of rarefaction (P≤0.05; Kruskal-Wallis one-way ANOVA) followed by women not on hormonal contraceptives, and those on oral contraceptives. (**D**) Sex workers on DMPA had significantly greater bacterial diversity in their vaginal microbiota than those not on hormonal contraceptives, at all depths of rarefaction (P≤0.05; Mann-Whitney U test). (**E**) Linear Discriminant Analysis effect size (LefSe) separated the vaginal microbiota of sex workers not on hormonal contraceptives versus those on DMPA based on enrichment of the bacterial genera listed, using 2.0 as a threshold for discriminative features, and P≤0.05 for statistical tests. DMPA: Depot-medroxyprogesterone acetate. Ns: Not significant. OCP: oral contraceptive pills. ^: Resolved to bacterial phylum. ^^: Resolved to bacterial class. *: P≤0.05.**: P≤0.01. Data is presented as mean ± SEM.

To identify taxa that significantly differed in the VMB of sex workers on DMPA as compared to NH, the LEfSe method was employed (26). Women on DMPA were more likely to be colonized by Operational Taxonomic Units (OTUs) from *Peptoniphilus, Finegoldia, Achromobacter, Verrucomicrobia, Varibaculum, Corynebacterium, Corynebacteriaceae, Verrucomicrobiaceae, Verrucomicrobiae, Akkermansia, Staphylococcus, Bacillales, Prevotella, Actinomycetales, Proteobacteria*, and *Actinobacteria*, while NH women were more likely to be colonized by OTUs within *Fermicutes*, and *Anaerococcus* (Figure 1E). Taken together, results suggest DMPA is positively associated with VMB diversity in healthy, asymptomatic Kenyan sex workers.

### Diversity correlates with activation of vaginal HIV-1 target cells in healthy, asymptomatic Kenyan sex workers

After determining that DMPA was associated with increased bacterial diversity in the vaginal tract, we sought to determine if there was a relationship between diversity of the VMB and vaginal cytokines and/or HIV-1 target cells in sex workers. Cytokines (IL-1α, IL-1ra, IL-1β, IL-8, IL-10, IL-17, sCD40L, IFN-γ, TNF-α, MIP-1α, MIP-1β, MIG-3, IP-10 and MCP-1) were quantified in the CVL (N=58), and linear regressions plotted. There were no significant correlations observed between diversity of the VMB (Shannon Diversity Index at 12744 reads) and any of the aforementioned cytokines.

Subsequently we sought to assess the relationship between VMB diversity and cervical HIV-1 target cells (CD4+CCR5+ T cells). There was no significant relationship between the percentage (Figure 2A; P=0.496, R^2^=0.009) or count (N) (Figure 2B; P=0.079, R^2^=0.06) of cervical CD4+CCR5+ T cells and VMB diversity (Shannon Diversity Index at 12744 reads). However, a significant positive correlation was observed between VMB diversity (Shannon Diversity Index at 12744 reads) and mean fluorescence intensity (MFI) of CCR5 expression on CD4+ T cells in the cervix (Figure 2C; P=0.009, R^2^=0.12), suggesting that with increasing bacterial diversity there are increased CCR5 receptors on cervical HIV-1 target cells (CD4+ cells). Thus, diversity of the VMB appears to correlate with activation of HIV-1 target cells in Kenyan sex workers with Nugent Scores <7. This provides a potential mechanism by which diversity of the VMB might influence inflammation in the female genital tract and impact susceptibility to HIV-1 in this cohort.

**Figure 2:**
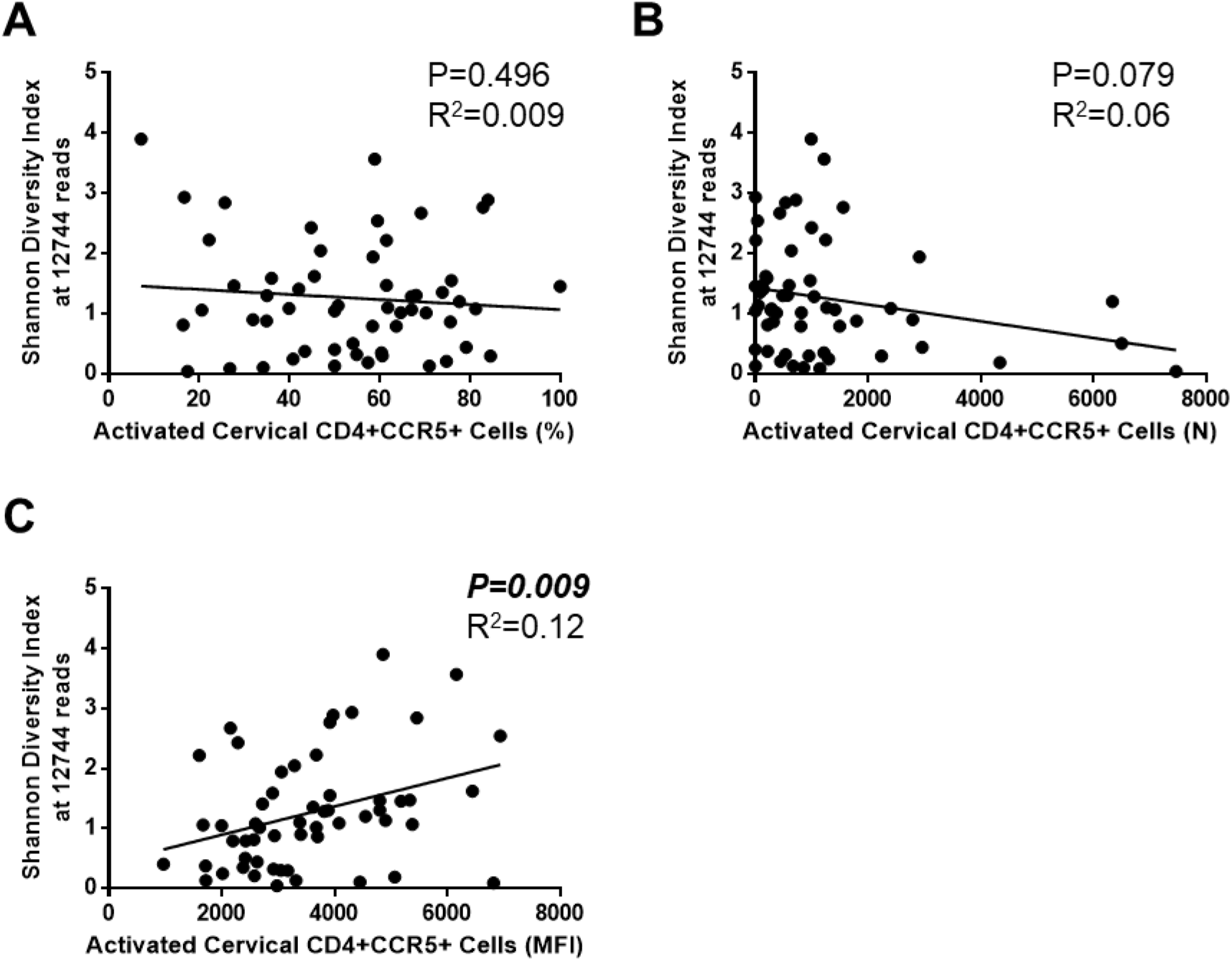
Diversity of the Vaginal Microbiota Correlates with Activated HIV-1 Target Cells, which might affect Susceptibility to HIV-1 in Sex Workers. Linear regression analyses were performed between diversity of the vaginal microbiota (Shannon Diversity Index at 12744 reads) and HIV-1 target cells (CD4+CCR5+) to determine if diversity of the vaginal microbiota correlated with factors that might impact susceptibility to HIV-1 in Kenyan sex workers (N=58). No significant relationship between the percentage **(A)** of cervical CD4+CCR5+ T cells and VMB diversity (Shannon Diversity Index at 12744 reads) was observed (P=0.496, R^2^=0.009), and no significant relationship between the number (count (N)) **(B)** of cervical CD4+CCR5+ T cells and VMB diversity (Shannon Diversity Index at 12744 reads) (P=0.079, R^2^=0.06) was observed. However, **(C)** a significant positive correlation was observed between VMB diversity (Shannon Diversity Index at 12744 reads) and activation status (MFI) of CD4+CCR5+ T cells in the cervix (P=0.009, R^2^=0.12), suggesting that with increasing bacterial diversity there is an increase in the CCR5 receptors on the HIV-1 target cells (CD4+ cells) present in the cervix. MFI: mean fluorescence intensity.

### DMPA does not alter the proportion of Lactobacillus-dominant VMBs in healthy, asymptomatic Kenyan sex workers

Typically a decreased risk of HIV-1 is associated with a low diversity, *Lactobacillus*-dominant VMB (11, 18, 27). Therefore, we examined whether hormonal contraceptives affected the proportion of sex workers with *Lactobacillus*-dominant VMB. The top 20 bacterial genera were plotted by method of contraception, as taxa bar charts (Figure 3A). The type of contraceptive did not significantly alter the proportion of sex workers with *Lactobacillus*-dominant VMB (>50% relative abundance). NH women were just as likely to have *Lactobacillus*-dominant VMB as those on DMPA (68%, 15/22 vs. 68%, 15/22, respectively), while those on OCPs had a slightly higher proportion of *Lactobacillus*-dominant women (79%, 11/14), but there was no significant difference in the proportion of *Lactobacillus-*dominant women between contraceptive groups (P=0.758; Chi-square). However, more sex workers on DMPA were borderline *Lactobacillus-*dominant and more genera were visually apparent in the taxa bar charts of these women, supporting our results demonstrating enhanced Shannon Diversity in women on DMPA (Figure 1C, D). Furthermore, the dominant species of *Lactobacillus* did not differ between groups (Table 1; P=0.737; Chi-square), and VMBs clustered by community state type (CST)(12), rather than method of contraception (Figure 3B, C) in principle coordinate analysis (PCoA) and cluster dendrograms. The gap statistic (Supplemental Figure 4A) indicated three clusters in the PCoA. PCoA ordination and Bray-Curtis dissimilarity distance were used to construct a heatmap, which also demonstrated clustering based on CSTs rather than type of hormonal contraceptive (Supplemental Figure 4B). Results suggest the overall proportion of Kenyan sex workers with Nugent scores <7 with *Lactobacillus*-dominant VMBs is not statistically significantly changed by DMPA. However there were more genera above 1% abundance in the taxa bar charts of sex workers on DMPA, supporting the observation of enhanced Shannon Diversity Index we observed in these women (Figure 1C, D).

**Figure 3:**
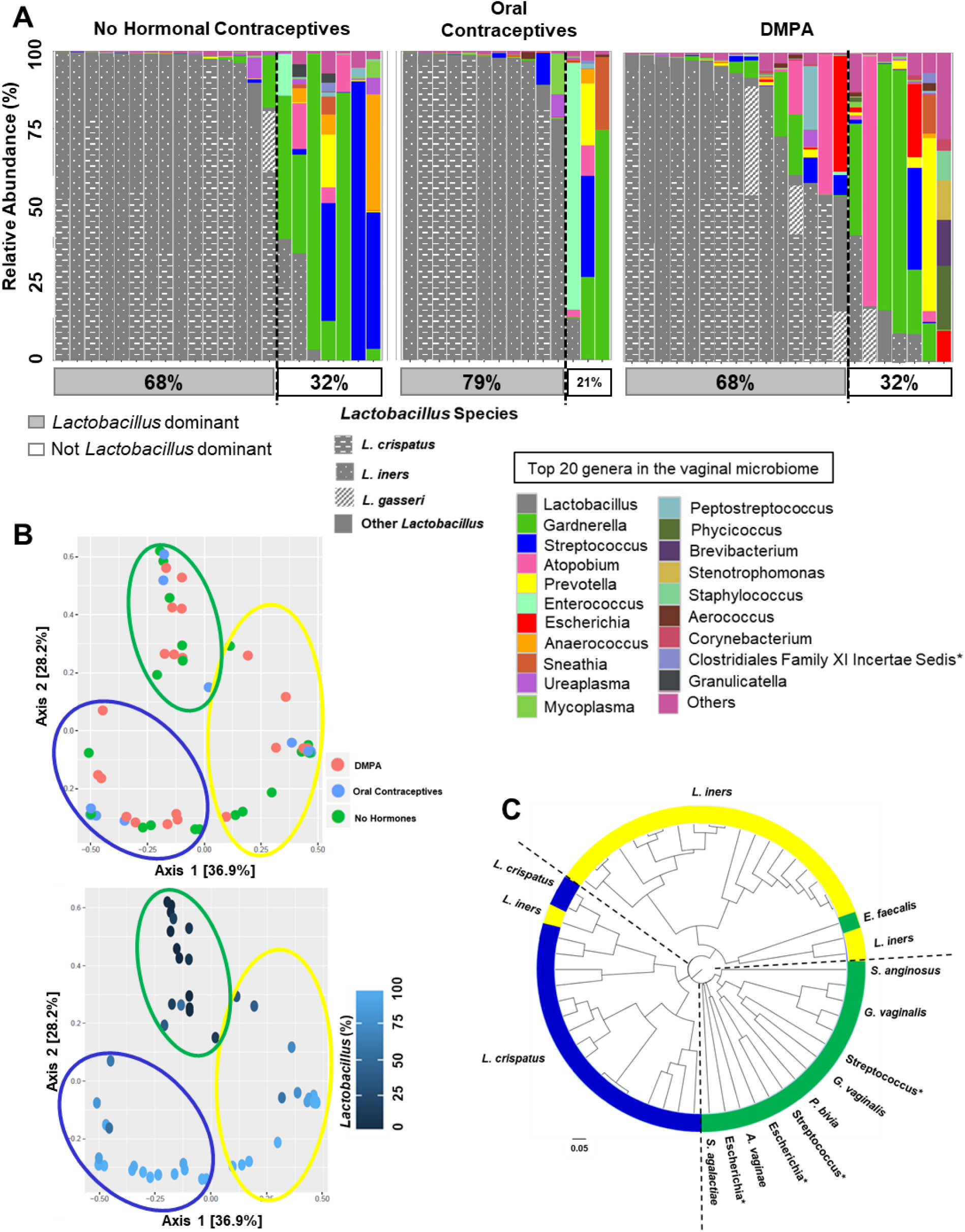
Contraceptives Did Not Affect *Lactobacillus* Dominance or Clustering. **(A**) The top 20 bacterial genera in the vaginal microbiota were plotted by relative abundance as taxa bar charts and compared between sex workers who were not on hormonal contraceptives (proliferative phase of the menstrual cycle, N=22), on oral contraceptives (N=14), or on DMPA (N=22). Each bar represents the vaginal microbiota of one woman. Each colour represents a different genus of bacteria, as indicated in the legend. Species of *Lactobacillus* are indicated in grey/patterns as per legend. Vaginal microbiota are ordered left to right in descending order of the relative abundance of *Lactobacillus*, and women to the left of the dashed lines are *Lactobacillus* dominant (>50% relative abundance, typical cut-point reported in the literature). The proportion of vaginal microbiota that were *Lactobacillus* dominant is listed as the percentage in the grey box below the bar chart. Sex workers not on hormonal contraceptives were just as likely to have a *Lactobacillus* dominant vaginal microbiota as those on DMPA, while those on oral contraceptives had a slightly higher proportion of women who were *Lactobacillus* dominant. *: Resolved to family level. (**B**) The principle coordinate analysis plots (PCoA) demonstrated the beta-diversity of the vaginal microbiota at the OTU level based on the Bray-Curtis dissimilarity matrix. The vaginal microbiota did not cluster by method of contraception, but clustered based on the community state types previously described (12); by the dominance of *Lactobacillus* in the vaginal microbiota. Sex workers dominated by *L. crispatus* (Community State Type (CST) I) are circled in blue, those dominated by *L. iners* (CST II) in yellow, and those women with highly diverse vaginal microbiota (CST IV) are circled in green. Axes = eigenvalues, a metric whose magnitude indicates the amount of variation captured in the PCoA axis. (**C**) A cluster dendrogram was calculated from Bray Curtis dissimilarities and used to visualize clustering of the vaginal microbiota. Three clusters were observed using the gap statistic (Supplemental Figure 4A), and in general, the previously described CSTs clustered together within the dendrogram. CSTI (blue): *L. cripatus* dominant, CSTIII (yellow): *L. iners* dominant, CSTIV (green): highly diverse. *: resolved to bacterial genus.

**Table 1:** Bacterial dominance in the vaginal microbiota of Kenyan sex workers does not differ by method of contraception.

|  | **>50% *L. crispatus*** | **>50% *L. iners*** | **Others** |
| --- | --- | --- | --- |
| No Hormonal Contraceptives | 6/22 (27%) | 9/22 (41%) | 7/22 (32%) |
| OCPs | 5/14 (36%) | 6/14 (43%) | 3/14 (21%) |
| DMPA | 7/22 (32%) | 6/22 (27%) | 9/22 (41%) |

P=0.737; Chi-square
DMPA: Depot-medroxyprogesterone acetate. *L.*: Lactobacillus. OCP: Oral contraceptive pills.
No Hormonal Contraceptives: proliferative phase of the menstrual cycle

### Healthy, asymptomatic Kenyan sex workers on DMPA have low vaginal glycogen and α-amylase

One factor believed to promote and maintain lactobacilli in the vaginal tract is glycogen, stored in vaginal epithelial cells, and available as free glycogen in vaginal fluid (13-15). Glycogen is a glucose polymer converted to disaccharides by α-amylase, and can be used by lactobacilli and other bacteria (16) as an energy source. Estrogen has been proposed to enhance glycogen in the human vaginal epithelium (28, 29), and glycogen is experimentally enhanced in the vaginal tissues of estrogen-treated hamsters, and non-human primates (30, 31). As DMPA is potently anti-estrogenic (4), we examined whether DMPA was associated with lower levels of free glycogen and α-amylase in the vaginal microenvironment as compared to NH women (with endogenous hormones), or those on estrogen-containing OCPs. As others have demonstrated the diverse vaginal microflora can overshadow the effect of hormones on vaginal glycosidases and lectins (22, 23), we restricted this analysis to women with Nugent scores ≤3. Vaginal glycogen (Figure 4A; N=16, 18; P=0.043) and α-amylase (Figure 4B; N=16, 16; P=0.0095) were significantly less abundant in the CVLs of sex workers on DMPA compared to NH women (2.04±0.42 vs. 5.98±1.72mg/mL; and 3.91±1.32 vs. 16.90±3.78mU/mL respectively). Furthermore, sex workers on estrogen-containing OCPs had the highest free vaginal glycogen, and significantly more vaginal glycogen than women on DMPA (Supplemental Figure 5A; N=13, 16; P=0.008), supporting the belief that estrogen is linked with enhanced vaginal glycogen.

**Figure 4:**
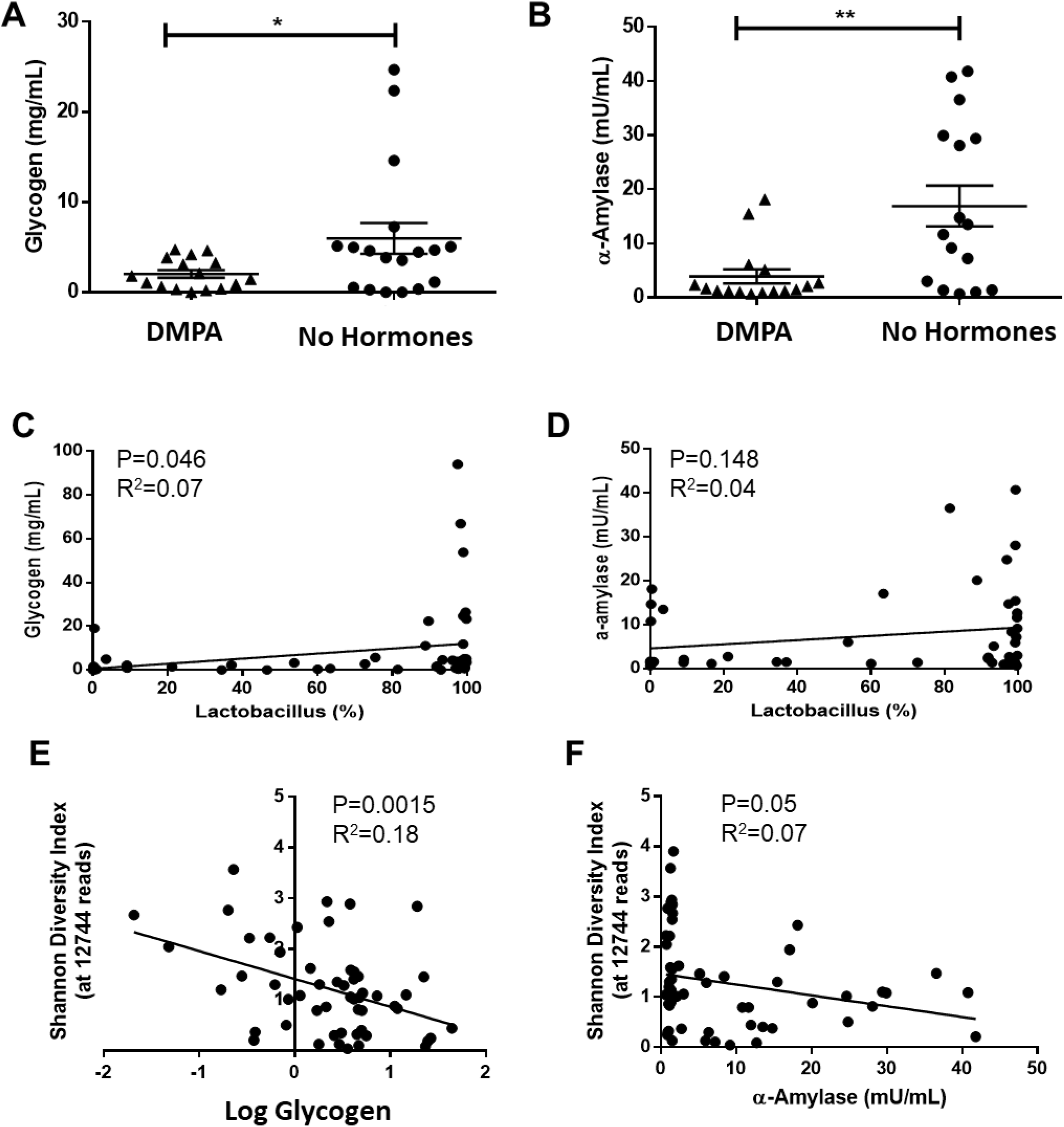
Sex Workers on DMPA have Low Vaginal Glycogen and α-Amylase. As the diverse vaginal microflora can strongly influence vaginal carbohydrates in CVLs and overshadow the effect of hormones on these types of parameters (22, 23), we restricted our analysis in (A) of vaginal glycogen and α-amylase to sex workers with Nugent scores ≤ 3. **(A)** Vaginal glycogen, a glucose polymer associated with vaginal colonization by *Lactobacillus* species, was significantly lower in the vaginal lavage of sex workers on DMPA (N=16) as compared to those not on hormonal contraceptives (proliferative phase of the menstrual cycle, N=18; Unpaired t test; P=0.043). **(B)** Similarly, α-amylase, the enzyme that breaks down glycogen into disaccharides which might act as substrates for vaginal lactobacilli, was significantly less abundant (Mann-Whitney; P=0.0095) in the vaginal lavage of sex workers on DMPA (N=16) as compared to sex workers who were not on hormonal contraceptives (proliferative phase of the menstrual cycle, N=16). **(C)** There was a positive correlation between free glycogen in the vaginal lavage and abundance of *Lactobacillus* species in the vaginal microbiota (N=54; Linear Regression; P=0.046, R^2^=0.07). **(D)** The relationship between α-amylase in the vaginal lavage and abundance of *Lactobacillus* species in the vaginal microbiota verged on significance (N=48; Linear Regression; P=0.148, R^2^=0.04). **(E)** A strong negative correlation between diversity of the vaginal microbiota (Shannon Diversity Index) and glycogen in the vaginal lavage was observed (N=54; Linear Regression; P=0.0015, R^2^=0.18). **(F)** There was also a negative correlation between bacterial diversity and α-amylase in the vaginal lavage (N=55; Linear Regression; P=0.05, R^2^=0.07). DMPA: Depot-medroxyprogesterone acetate.

As glycogen is believed to maintain lactobacilli in the vaginal tract we assessed the relationship between free glycogen and *Lactobacillus* abundance. Free glycogen in the CVL positively correlated with abundance of *Lactobacillus* species in the VMB (Figure 4C; N=54; Linear Regression; P=0.046, R^2^=0.07), while the relationship between α-amylase and *Lactobacillus* verged on significance (Figure 4D; N=48; Linear Regression; P=0.148, R^2^=0.04). Additionally, we observed a strong negative correlation between bacterial diversity and vaginal glycogen (Figure 4E; N=54; P=0.0015, R^2^=0.18), and a negative correlation between bacterial diversity and α-amylase (Figure 4F; N=55; P=0.05, R^2^=0.07). Together, results suggest DMPA has a negative impact on vaginal glycogen and α-amylase in sex workers, two factors thought to be important for vaginal colonization by lactobacilli (13-15, 32).

### DMPA is associated with vaginal microbial diversity in humanized mice

Given our stringent recruitment criteria and consequently high attrition rate, our sample size was limited (Supplementary Figure 1). Therefore, we decided to experimentally validate the main effects of DMPA on VMB diversity and vaginal glycogen. We recently optimized a humanized mouse model, reconstituted with human immune cells, that demonstrates HIV-1 infection following intravaginal challenge (33). We examined the effect of DMPA on diversity of the VMB in this model. Since humanized mice have disrupted/irregular reproductive cycles due to radiation treatment administered to deplete bone marrow (required for engraftment of human immune cells), we treated humanized mice with physiological concentrations of estradiol (E2) or DMPA for 14 days, and assessed alpha-diversity from vaginal washes using the same metrics we had in women. To ensure subcutaneous injection of DMPA could induce circulating concentrations similar to women, we determined MPA levels in the serum of mice injected with different concentrations of DMPA, as described in Methods. Our results indicated that 2mg of subcutaneous DMPA consistently led to concentrations of MPA in humanized mice that were similar to those seen in women (Supplemental Figure 2A, D). As in the sex worker cohort, there were no significant differences in observed species (Figure 5A), or Chao1 (Figure 5B), in the VMB of humanized mice given E2 versus DMPA. However, the Shannon Diversity Index was significantly greater in DMPA-treated humanized mice compared with E2 (Figure 5C). LEfSe (26) identified taxa that significantly differed between humanized mice on DMPA versus E2. Humanized mice on DMPA were significantly more likely to be colonized by 30 different taxa than those on E2 (Figure 5D). Although bacterial species observed in the vaginal tract of humanized mice (Supplemental Figure 6) were different from those in women, results recapitulated our clinical observations in sex workers; that DMPA is associated with diversity of the humanized mouse VMB.

**Figure 5:**
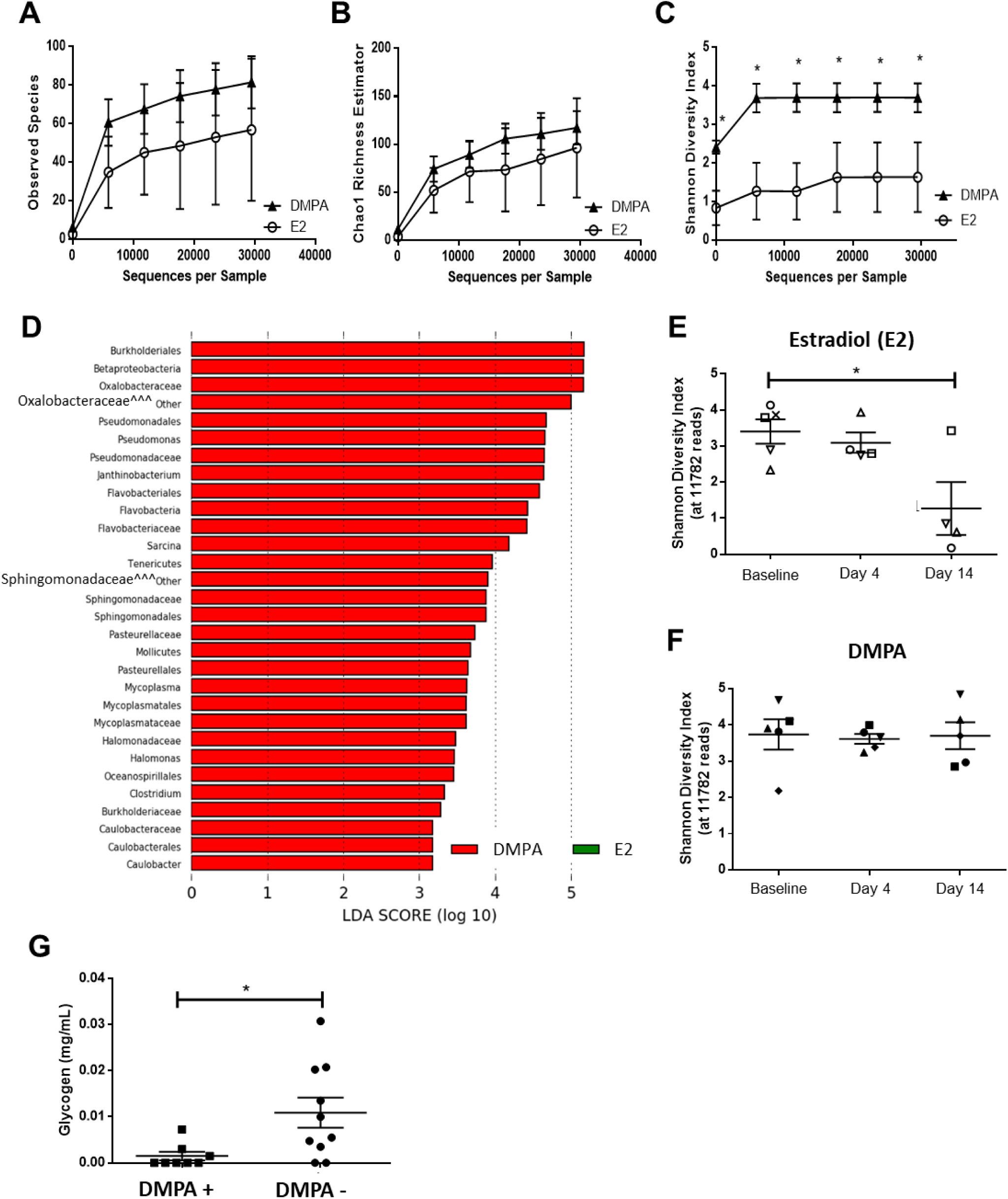
DMPA is associated with Diversity of the Vaginal Microbiota, as a Result of Hypo-estrogenism, and Suppresses Vaginal Glycogen in Humanized Mice. Bacterial richness and diversity were quantified in the vaginal microbiota of humanized mice treated with estradiol (E2, N=4) or DMPA (N=5) for 14 days. **(A)** No significant difference in Observed Species was seen between humanized mice on E2 versus DMPA. (**B**) Similarly, there was no significant difference in bacterial richness (estimated by Chao1) between humanized mice on E2 versus DMPA. (**C**) However, mirroring our clinical data, humanized mice on DMPA had significantly greater bacterial diversity in their vaginal microbiota than those on E2, at all depths of rarefaction (P≤0.05). (**D**) Linear Discriminant Analysis effect size (LefSe) separated the vaginal microbiota of humanized mice treated with E2 versus DMPA based on enrichment of the bacterial genera listed, using 2.0 as a threshold for discriminative features, and P≤0.05 for statistical tests. **(E)** The temporal effect of E2 and DMPA on diversity of the humanized mouse vaginal microbiota was assessed over time. Humanized mice treated with estradiol had a significant reduction in diversity of the vaginal microbiota, as assessed by the Shannon Diversity Index (N=4; P=0.02) over time. Each humanized mouse is denoted by a unique symbol in the graphs to allow for comparisons in bacterial diversity between days, and for comparisons with taxa bar charts (Supplemental Figure 6A). **(F)** Diversity of the vaginal microbiota in humanized mice treated with DMPA remained chronically high (N=5; P=0.96), suggesting DMPA does not directly increase bacterial diversity, but rather that the presence of estradiol reduces bacterial diversity in the vaginal microbiota of humanized mice. DMPA induces hypo-estrogenism in women (4) (Supplemental Figure 2A), it thus stands to reason that in the absence of endogenous estrogen diversity of the vaginal microbiota is not suppressed, and thus DMPA correlates with diversity. Each humanized mouse is denoted by a unique symbol in the graphs to allow for comparisons in bacterial diversity between days, and for comparisons with taxa bar charts (Supplemental Figure 6B). **(G)** Glycogen was quantified in the vaginal homogenate of humanized mice (controls, no hormonal treatment, N=10) and those treated with DMPA (N=8). Vaginal glycogen was significantly lower in the humanized mice treated with DMPA (Mann-Whitney U test; P=0.017). E2: Estradiol. DMPA: Depot-medroxyprogesterone acetate. ^^^: Resolved to bacterial family. *: P≤0.05. Data is presented as mean ± SEM.

### Vaginal microbial diversity is minimized by estradiol in humanized mice

We collected vaginal lavage from humanized mice at baseline and during hormone treatment (day 4, 14), and assessed the temporal effect of E2 and DMPA on diversity of the humanized mouse VMB. Diversity of the VMB was significantly reduced over time, as assessed by the Shannon Diversity Index (Figure 5E; N=4; P=0.02), and taxa bar charts (Supplemental Figure 6A) in E2-treated humanized mice. Conversely, VMB diversity remained consistently high in DMPA-treated humanized mice (Figure 5F; N=5; P=0.96) (Supplemental Figure 6B). This suggests estradiol may be important in reducing and maintaining low bacterial diversity. As DMPA induces hypo-estrogenism (4) (Supplemental Figure 2A), it stands to reason that bacterial diversity is not suppressed in the absence of estrogen, and thus DMPA correlates with VMB diversity.

Although bacterial diversity is not significantly changed over time in DMPA-treated humanized mice, the relative abundance of bacterial genera does appear to shift in response DMPA-treatment (Supplemental Figure 6B), which could potentially alter immune cell populations in the vaginal tract, and impact HIV-1 susceptibility.

### DMPA is associated with low vaginal glycogen in humanized mice

Since estradiol is thought to enhance glycogen in vaginal epithelium, and DMPA is hypo-estrogenic, we posited we would see similar effects of DMPA on glycogen in humanized mice, as we had seen in the Kenyan sex worker cohort. Glycogen was quantified in vaginal homogenates from humanized mice that had received DMPA or not (no hormonal treatment). Vaginal glycogen was significantly lower in DMPA-treated humanized mice compared with control untreated mice (Figure 5G; 1.5×10^-3^±9.0×10^-4^vs.1.1×10^-2^±3.0x×10^-3^mg/mL; N=10, 8; P=0.017). Thus, DMPA suppresses vaginal glycogen in humanized mice, similar to the correlation we observed in sex workers.

### DMPA enhances susceptibility to HIV-1 in humanized mice

DMPA is associated with increased susceptibility to HIV-1 in women (1), and women with high diversity VMB are more susceptible to HIV-1 (11). Here, we demonstrate DMPA is associated with VMB diversity in Kenyan sex workers and humanized mice, which may contribute to increased HIV-1 risk. No studies have demonstrated a direct link between DMPA and enhanced susceptibility to HIV-1 in humanized mice. We thus experimentally assessed whether DMPA-treated humanized mice were more susceptible to HIV-1 in a heterosexual model of viral transmission (33) compared to untreated, control humanized mice. Humanized mice (control, no hormonal treatment, N=20) and DMPA-treated humanized mice (N=22) were challenged intravaginally with 10^5^ TCID50/mL NL4.3-Bal-Env HIV-1 (33, 34). HIV-1 viral load in peripheral blood was quantified by clinical RT-PCR 3-5 weeks following challenge (33). The proportion of infected DMPA-treated humanized mice was 17/22 (77%), which was significantly higher (P=0.014) than those not receiving DMPA (7/20, 35%) (Figure 6). These results show in an experimental model, treatment with DMPA significantly enhanced HIV-1 infection following intravaginal viral exposure.

**Figure 6:**
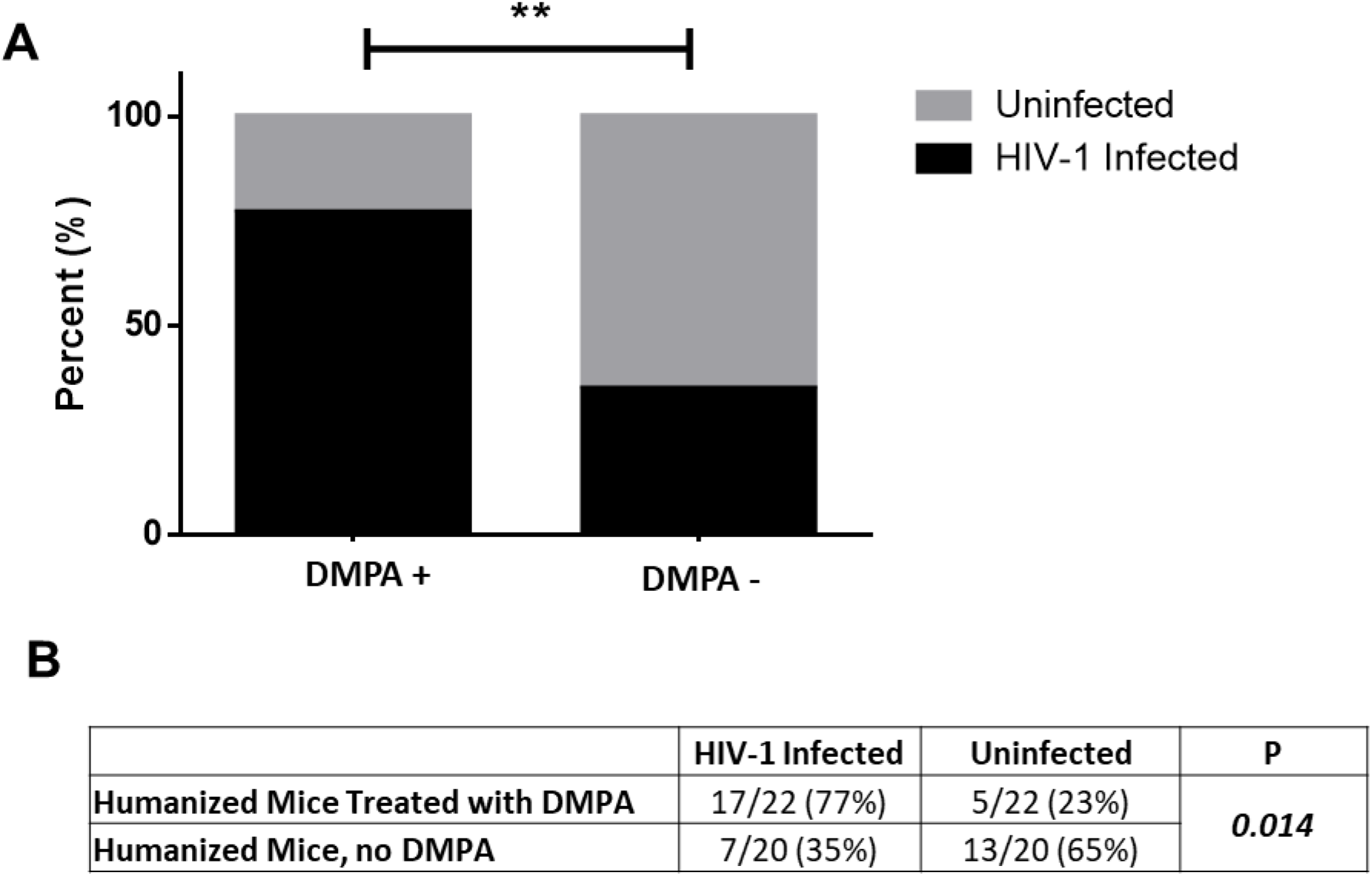
DMPA Increases Susceptibility to HIV-1 in Humanized Mice. In order to determine if humanized mice treated with physiological levels of DMPA were at a greater risk of contracting HIV-1 following intravaginal challenge (model of heterosexual transmission), control humanized mice (no hormonal treatment, N=20) and humanized mice given 2mg DMPA (N=22) were challenged intravaginally with HIV-1. **(A)** The infection rate in humanized mice that had been given DMPA (77%) was higher than the infection rate observed in untreated control humanized mice (35%) (Chi-Square; P=0.014). **(B)** The proportion of humanized mice that became infected following intravaginal challenge with HIV-1 was significantly greater (Chi-Square; P=0.014) than the proportion of untreated control humanized mice that became infected following intravaginal challenge. Humanized mice were considered to be infected if there was a detectable HIV-1 viral load in the peripheral blood, as quantified by clinical RT-PCR by 3 or 5 weeks following viral challenge. DMPA: Depot-medroxyprogesterone acetate.

## Discussion

This study provides compelling clinical evidence linking vaginal bacterial diversity to use of the hormonal contraceptive DMPA in healthy, asymptomatic Kenyan sex workers. We also demonstrate VMB diversity correlates with T cell activation, and that vaginal glycogen and α-amylase are low in sex workers on DMPA. We recapitulated results in an experimental model, where DMPA-treated humanized mice had greater VMB diversity and less vaginal glycogen. These mice also had enhanced susceptibility to HIV-1 following intravaginal challenge. Based on these results we posit a potential mechanistic model linking DMPA-induced hypo-estrogenism to changes in the VMB and microenvironment that might impact susceptibility to HIV-1 (Figure 7). These results represent substantial progress towards understanding the biological link between MPA, the vaginal microenvironment, and susceptibility to HIV-1 in sex workers without clinical BV.

**Figure 7:**
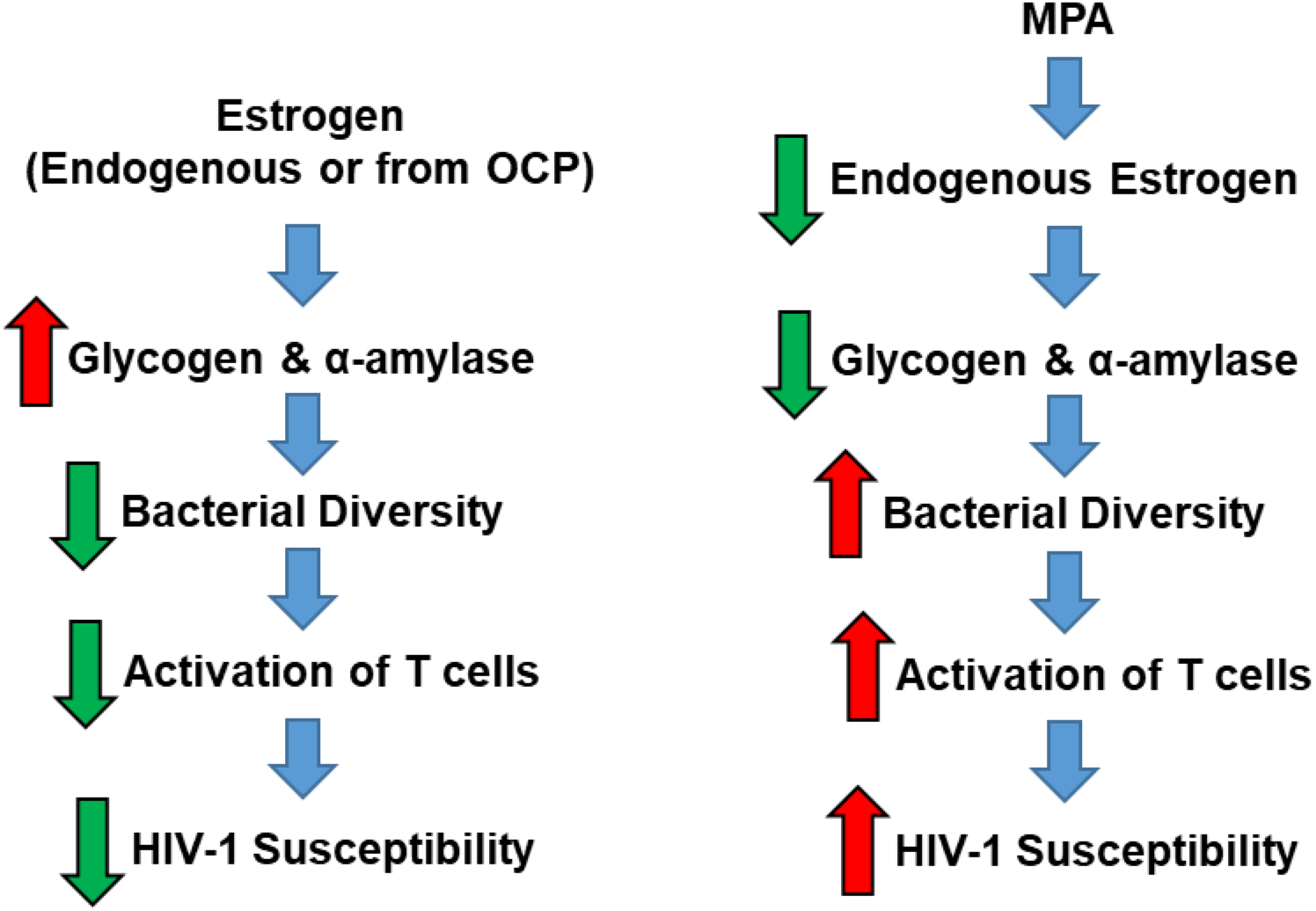
Proposed Mechanism linking DMPA to changes in Vaginal Microenvironment and HIV-1 Susceptibility in Kenyan Sex Workers. **(A)** Taken together our results suggest that estrogen (either from endogenous sources or oral contraceptive pills) is associated with greater quantities of vaginal glycogen and α-amylase, and minimal bacterial diversity within the vaginal microbiota of sex workers. This stable, uniform microbiota is not associated with inflammatory cytokines or activated T cells, and as a result is not associated with susceptibility to HIV-1. In contrast, the hypo-estrogenism resulting from use of MPA lowers key metabolic products (ie. glycogen, and α-amylase) that can be used by certain protective bacterial species, and the change in substrates allows other bacteria to colonize the vaginal microbiota, in effect increasing bacterial diversity. Diversity of the vaginal microbiota was positively correlated with activation of HIV-1 target cells in the vaginal tract, indicating that the subtle shift in bacterial diversity might be sufficient to attract immune attention, and thus contribute to enhanced susceptibility to HIV-1 in sex workers. OCP: oral contraceptive pills. MPA: Medroxyprogesterone acetate (active ingredient in DMPA).

Low diversity *Lactobacillus*-dominated VMB are thought to protect against HIV-1 in women (11, 35). Lactobacilli appear to provide non-specific defense against a broad range of pathogens via production of lactic acid, hydrogen peroxide, and anti-microbial bacteriocins, and by providing a physical/neutralizing barrier that inhibits other bacteria/pathogens (4). The current understanding of the relationship between vaginal bacterial diversity and HIV-1 susceptibility stems from the BV literature, where women with BV (high diversity VMB) are more prone to acquiring HIV-1 than those without (36-38); likely a result of increased HIV-1 target cells attracted by the inflammation associated with the polymicrobial microbiota. However, it is increasingly clear that bacterial diversity, *even in the absence of clinical BV* might confer greater susceptibility to HIV-1 (11, 37), as women with intermediate vaginal flora (intermediate diversity) still had a 1.5X increased risk of HIV-1 acquisition in a meta-analysis (37). Here, we examined the effect of MPA on VMB diversity in Kenyan sex workers, excluding women with BV. Importantly, we found the VMB of sex workers with Nugent scores <7 who were on DMPA was significantly more diverse than those with Nugent scores <7 who were not on hormonal contraceptives. This relationship was still apparent when we excluded sex workers with intermediate vaginal flora (Supplemental Figure 3), suggesting MPA is associated with VMB diversity in Kenyan sex workers with low Nugent Scores, without BV. The relationship between DMPA and bacterial diversity is important because women with diverse VMBs had a 4-fold increased risk of HIV-1 in a recent study (11), and we found diversity correlated with activation of vaginal CD4+ HIV-1 target cells in the Kenyan sex worker cohort. Although BV, which is an inherently high diversity VMB, is linked to increased risk of HIV-1 (36) it is important to consider that women with BV are not the only women acquiring HIV-1; women with intermediate vaginal flora (intermediate diversity) still have a 1.5X increased risk of HIV-1 acquisition (37). Additionally, meta-analyses demonstrate DMPA is independently associated with a 1.4X increased risk of HIV-1 (1). Therefore, our results connecting DMPA to diversity of the VMB in Kenyan sex workers without BV provide novel insight into the mechanisms by which healthy, asymptomatic sex workers using DMPA might be more prone to acquiring HIV-1 than those not taking hormonal contraceptives.

Although we and others (10) demonstrate that DMPA is associated with enhanced diversity of the VMB by 16S rRNA gene sequencing in women with Nugent scores <7, it is important to note that hormonal contraceptives (DMPA and OCPs) are typically associated with a reduced incidence and prevalence of BV, as assessed by Nugent scoring (39). The implication is that DMPA and OCPs therefore reduce “bacterial diversity” but the bacterial diversity assessed by Nugent scoring is based on microscopic visualization of bacterial morphotypes and may not be as sensitive in assessing bacterial diversity as the α-diversity metrics that can be calculated by 16S rRNA gene sequencing. In the present study, when we directly compared bacterial diversity by 16S rRNA sequencing in Kenyan sex workers without BV those on DMPA were significantly more diverse than those who were not on hormonal contraceptives. Thus, given that systematic reviews find a reduced rate of BV in women on DMPA and OCPs (39), the relationship between bacterial diversity and DMPA may not have been as obvious if our study had included women with BV. However, because BV is a clinical condition reported to overshadow the effect of hormones on vaginal biomarkers of microbial health (glycosidases, lectins) in CVLs (22, 23), and because we already know BV is associated with increased risk of HIV-1 (36) we chose to exclude women with Nugent scores 7-10 and focus on understanding how DMPA and diversity of the VMB in the absence of BV might influence HIV-1 susceptibility in Kenyan sex workers.

To confirm the DMPA/diversity relationship observed in our cohort of Kenyan sex workers, we treated humanized mice with DMPA and found MPA also associated with VMB diversity in humanized mice. However further examination suggested that in humanized mice DMPA did not increase bacterial diversity, but rather estradiol treatment suppressed diversity, rendering the humanized mouse VMB similar to the typically mono-colonized VMB of women. Importantly, the VMB of women and mice are different. Women often have a low diversity VMB dominated by one bacterial species (usually lactobacilli) whereas mice have high diversity VMB, not dominated by any bacterial species. It is possible mice carry the maximal amount of VMB diversity, and thus bacterial diversity can only be minimized. Conversely in women there may be room for increased or decreased diversity. Thus, DMPA may directly or indirectly increase diversity of the VMB in women, but be unable to do so in maximally diverse humanized mice. Either way, determining estradiol suppresses VMB diversity in humanized mice suggests DMPA affects bacterial diversity in Kenyan sex workers via its hypo-estrogenic effects (Supplemental Figure 2A).

We also found lower vaginal glycogen and α-amylase in Kenyan sex workers on DMPA. Glycogen is a glucose polymer that can be converted to disaccharides by α-amylase, and utilized by bacteria like lactobacilli as an energy source (16). These two factors are thought to select for a *Lactobacillus-*dominated microbiota in the human vagina, which is protective against many STIs (13-15, 32). A change in carbohydrate resources could negatively impact protective bacteria, allow other species not as reliant on glycogen to colonize the vaginal tract, and enhance bacterial diversity. This may be particularly true in a sex worker cohort, where women are more likely to be exposed to a variety of bacteria because they likely have more sex, multiple partners, and different sexual practices than women who are not sex workers. Importantly, MPA was also associated with low vaginal glycogen in humanized mice, even though *Lactobacillus* is not dominant in the mouse vagina. This suggests that regardless of host species, MPA affects carbohydrate resources within the vaginal microenvironment, and bacterial species able to access the vaginal tract and thrive under the new conditions respond accordingly. Additionally, estrogen is believed to enhance glycogen in human vaginal epithelium (28, 29), and experimentally enhances vaginal glycogen in hamsters, and non-human primates (30, 31), via incompletely understood mechanisms. DMPA is hypo-estrogenic (Supplemental Figure 2A) and might therefore restrict glycogen by reducing endogenous estrogen. The effect of MPA on estrogen-mediated pathways supporting glycogen in the vaginal tract may not be host species specific, which would explain dampened vaginal glycogen in DMPA-treated sex workers and humanized mice. Therefore, our findings in two different species suggest MPA changes bacterial resources in the vaginal tract, and supports a diverse array of bacteria under the altered conditions.

Although DMPA is associated with increased bacterial diversity, low vaginal glycogen, α-amylase and activated CD4+ target cells in the vaginal tract of Kenyan sex workers, and some of these results were recapitulated in the humanized mice, we wanted to determine if MPA enhanced HIV-1 susceptibility in these mice. We challenged humanized mice intravaginally with HIV-1 and DMPA-treated mice were more likely to become infected than controls (77% vs. 35%, P=0.014). Although MPA could enhance HIV-1 infection via other mechanisms, results suggest MPA might alter the humanized mouse VMB by slightly shifting the relative abundance of bacterial genera (Supplemental Figure, 6B). Lowering key carbohydrate resources like glycogen could select for other bacterial species, changing their relative abundance, and the shift in bacteria thus attracts immune attention, and activation of HIV-1 target cells in the vagina leading to greater HIV-1 susceptibility. To further support this hypothesis, we found a positive correlation between activation of cervical CD4+ cells and diversity of the VMB in our cohort of Kenyan sex workers, while others demonstrated activated cervical HIV-1 target cells (CD4+CCR5+CD25+) were increased in women on long-term progestin-only contraceptives (DMPA, norethisterone enanthate) (2), and a 17-fold increase in activated cervical target cells (CD4+CCR5+CD38+HLA-DR+) in women with high diversity VMB (11) was reported. Furthermore, women with high diversity VMB have elevated inflammatory cytokines (IL-1β, IL-1α, and IL-8) (40), and vaginal bacteria affect inflammatory responses and barrier function in the female reproductive tract (41-44). In fact, diverse VMB and use of hormonal contraceptives (all contraceptives combined, mainly progestins) were independently identified as the strongest predictors of genital inflammation (44), which is a key determinant of HIV-1 risk. There is also evidence DMPA enhances susceptibility to SIV in non-human primates via inflammatory pathways (45).

Several studies report the effect of hormonal contraceptives on the VMB (2, 5-10, 42-44). While some studies found subtle VMB shifts in women on hormonal contraceptives like enhanced *Lactobacillus* species (5, 8), reduced total bacteria (9), *G. vaginalis* (9), or lactobacilli (6), the majority do not show any major effects of hormonal contraceptives on VMB clustering (2, 7, 43) or associations between hormonal contraceptives and the proportion *Lactobacillus*-dominant VMB (42). Similarly, the taxa bar charts, PCoAs, heatmap, and cluster dendrogram in our study do not demonstrate any major effects of hormonal contraceptives on VMB clustering or associations between hormonal contraceptives and the proportion *Lactobacillus*-dominant VMB (except more borderline *Lactobacillus*-dominance in DMPA group), suggesting the effect of MPA on the VMB is more subtle than initially expected. However, our study provides a comprehensive assessment of MPA on VMB diversity in Kenyan sex workers, excluding BV as a confounder. Most previous studies employed targeted methods (qPCR (9), bacterial culture (5, 6), microarrays (7)) to draw conclusions which could bias results. Further, confounding variables including ethnicity, method of contraception (ie. all forms or progestins together), varying lengths of time on hormonal contraceptive, and BV were major potential confounders in these studies, making it difficult to discern the effect of hormonal contraceptives on the VMB. The exclusion of women with BV is important considering recent reports demonstrating its strong influence over the effect of hormones on vaginal parameters (22, 23), and positive correlation with microbial diversity (20). Here, we minimized potential biases by conducting untargeted 16S rRNA gene sequencing of the VMB in sex workers of the same ethnicity, controlling for recent sex (abstinence) and length of time on hormonal contraceptive (>6 months and within 3-4 weeks of most recent injection) in our inclusion criteria, not grouping progestins, and excluding sex workers with BV.

This study demonstrates an important association between DMPA and diversity of the VMB in Kenyan sex workers, and adds to the literature suggesting no major effect of hormonal contraceptives on VMB clustering (2, 7, 43) or association between hormonal contraceptives and the proportion of *Lactobacillus*-dominant women (42). It is however important to note that a shift in the VMB by 1% may have major biological consequences, considering the quantity of bacteria in the vaginal tract is 1010 or higher (9). We are not however the only group to examine the effect of DMPA on VMB diversity. A study by Birse et al., 2017 found no significant association between DMPA and bacterial diversity (Shannon Diversity Index) in non-sex workers, however women with BV were not excluded (43). The results of the present study support those of Jespers et al., 2017 who demonstrated that injectable progestins (including DMPA) were associated with VMB diversity and reduced concentration of lactobacilli in African women (10). Although another study included non-sex workers of varying ethnicities, and included BV, bacterial diversity (by inverse Simpson’s index) was significantly lower in women on OCPs than DMPA or not on hormonal contraceptives, and although not statistically significant, women on DMPA had greater bacterial diversity than women not on hormonal contraceptives (8). These studies support the association between MPA and VMB diversity in healthy, asymptomatic Kenyan sex workers that was observed in our study which may have been seen because of our strict inclusion/exclusion criteria, particularly the exclusion of women with BV.

Our study was not without limitations. Factors including ethnicity, cultural background, and vaginal douching affect the VMB (12, 18-20, 46-48). We therefore recruited sex workers of the same ethnicity, and geographical region to minimize confounders. Although we collected information on vaginal douching, we do not have information on intravaginal drying practices, which are associated with VMB diversity (43). However, because we did not see a significant difference in the proportion of sex workers practicing vaginal douching, we do not anticipate a difference in women performing vaginal drying, by method of contraception. One limitation of our study was the relatively small clinical study sample size. This was due to our strict inclusion/exclusion criteria. However, these same criteria were likely key in our ability to demonstrate the subtleties of the effect of MPA on the sex worker VMB. To compensate for small size we experimentally recapitulated clinical study results in humanized mice. In the present study, we collected samples from NH women on days 5-10 of the menstrual cycle to minimize cycle variability and standardize cycle phase. However this phase is the estrogen high phase of the menstrual cycle, and because we did not recruit sex workers during the progesterone high phase, one could question if differences in diversity were due to sample collection time for NH women. However two independent prospective studies have shown the Shannon Diversity Index remains stable across the menstrual cycle (19, 48). We thus believe results represent an accurate depiction of the effect of MPA on VMB diversity and the vaginal microenvironment in sex workers, especially given results of recent studies (8, 10).

The vaginal tract represents a major site of HIV-1 acquisition in women, and use of DMPA is consistently associated with enhanced HIV-1 susceptibility. As bacteria lining the vaginal tract regulate genital inflammation, and thus recruitment of HIV-1 target cells, understanding effects of MPA on the VMB of sex workers is an important public health issue. This is especially true considering DMPA is used by 8M women in sub-Saharan Africa (3), where HIV-1 is endemic. Herein, we demonstrate MPA is associated with VMB diversity, and suppression of vaginal glycogen and α-amylase in Kenyan sex workers; two factors believed to promote vaginal colonization by protective bacteria. We also show MPA has similar effects in humanized mice, and enhances susceptibility to HIV-1 following intravaginal challenge. We thus propose one mechanism by which MPA increases susceptibility to HIV-1 in sex workers is via suppressing endogenous estrogen, which is responsible for maintaining glycogen and α-amylase, factors thought to select for protective bacteria. As a result of the change in substrates other species not as reliant on these substrates access and compete for space and colonize the vaginal tract. The subsequent increase in bacterial diversity might activate HIV-1 target cells in the vaginal tract, and enhance susceptibility to HIV-1 in Kenyan sex workers. Future studies aimed at exploring MPA-mediated mechanisms of HIV-1 susceptibility in women, especially non-sex workers, and humanized mice are warranted.

## Methods

### Experimental design

This prospective cross-sectional study was approved by McMaster University’s Hamilton Integrated Research Ethics Board (HiREB 0332-T), University of Manitoba (B2015:033) and Nairobi/Kenyatta National Hospital (KNH-ERC P132/03/2015). Our objectives were to determine if DMPA affected 1) VMB diversity, 2) the proportion of *Lactobacillus-*dominant sex workers, 3) VMB clustering, and 4) vaginal glycogen and α-amylase. Results were replicated in humanized mice, where objectives were to experimentally confirm DMPA affected 1) VMB diversity, 2) vaginal glycogen, and 3) susceptibility to HIV-1.

Study participants were screened/enrolled through the Sex Workers Outreach Programme (SWOP), Pumwani Sex Worker cohort in Nairobi, Kenya (49). Women were recruited January 2015 to April 2016, following study outline (Supplemental Figure 1), and strict inclusion/exclusion criteria below. Written informed consent, and demographic information (Table 2) was obtained following explanation of study and potential risks.

**Table 2:** Demographic information for study participants.

|  | No Hormone<br>N=22 | OCP<br>N=14 | DMPA<br>N=22 | P-value |
| --- | --- | --- | --- | --- |
| **Mean Age (Range), Years<sup>a</sup>** | 29.9 (19-39) | 31.1 (24-45) | 27.8 (19-39) | 0.190 |
| **Years of Education, Mean (SD)<sup>a</sup>** | 10.5 (2.0) | 11.4 (2.3) | 10.1 (3.0) | 0.294 |
| **Marital Status<sup>b</sup>** |  |  |  |  |
| Married or Steady Partner | 8 (36%) | 7 (50%) | 9 (41%) | 0.775 |
| Unmarried | 13 (59%) | 7 (50%) | 13 (59%) |  |
| Unknown | 1 (5%) | 0 (0%) | 0 (0%) |  |
| **Menstrual Cycle Stage<sup>b</sup>** |  |  |  |  |
| Proliferative | 21 (95%) | 0 (0%) | 0 (0%) | <0.001 |
| Secretory | 1 (5%) | 0 (0%) | 0 (0%) |  |
| Hormonal Contraceptive | 0 (0%) | 14 (100%) | 22 (100%) |  |
| **Contraception, Parity** |  |  |  |  |
| Length of Time on Contraception, Mean (SD), Months<sup>c</sup> | N/A | 20.6 (11.3) | 37.4 (31.7) | 0.241 |
| Time Since Last DMPA Injection, Mean (SD), Days | N/A | N/A | 33.1 (15.1) |  |
| Number of Pregnancies, Mean (SD)<sup>d</sup> | 1.9 (1.2) | 1.7 (0.8) | 1.7 (0.8) | 0.894 |
| **Substance Use** |  |  |  |  |
| Alcohol Use<sup>b</sup> | 10 (45%) | 6 (43%) | 12 (55%) | 0.780 |
| Drug Use<sup>b</sup> | 0 (0%) | 1 (7%) | 1 (5%) | 0.508 |
| **Sex Work** |  |  |  |  |
| Duration of Sex Work, Mean (SD), Years<sup>a</sup> | 2.1 (0.9) | 2.5 (1.3) | 1.9 (0.8) | 0.442 |
| Number of Clients in the Last 5 Days, Mean (SD)<sup>d</sup> | 3.4 (5.6) | 0.9 (1.5) | 1.6 (2.2) | 0.525 |
| Number of Regular Partners, Mean (SD)<sup>d</sup> | 4.6 (3.6) | 3.7 (2.3) | 3.9 (4.6) | 0.510 |
| Sex without Condom in Last 5 Days<sup>b</sup> | 2 (9%) | 2 (14%) | 1 (5%) | 0.646 |
| Sex without Condom in Last 24 Hours | 0 (0%) | 0 (0%) | 0 (0%) |  |
| **Vaginal Douching** |  |  |  |  |
| Last 24 Hours<sup>b</sup> | 6 (27%) | 1 (7%) | 4 (18%) | 0.240 |
| Last 3 Days<sup>b</sup> | 7 (32%) | 1 (7%) | 4 (18%) | 0.174 |
| **Nugent Score (%)<sup>b</sup>** |  |  |  |  |
| 0-3 | 18 (82%) | 13 (93%) | 17 (77%) | 0.487 |
| 4-6 | 4 (18%) | 1 (7%) | 5 (23%) |  |
| a. ANOVA |  |  |  |  |
| b. Chi-square |  |  |  |  |
| c. Mann-Whitney U test |  |  |  |  |
| d. Kruskal-Wallis ANOVA |  |  |  |  |

DMPA: Depot-medroxyprogesterone acetate. OCP: Oral contraceptive pills. SD: Standard deviation. No Hormone: proliferative phase of the menstrual cycle

Women were included if >18 years old, pre-menopausal, willing to undergo pelvic exams, intact uterus and cervix, in good health and negative for STIs (gonorrhea, chlamydia, *Trichomonas vaginalis*, syphilis, HIV), negative for yeast infection, Nugent Score <7 at screening, could abstain from douching 24 hours, and sexual activities 12 hours before sample collection, and use condoms for 36 hours prior to abstinence, and had not been in sex work >5 years. Sex workers on DMPA were using injectable DMPA >6 months and within 3-4 weeks of most recent injection. Sex workers on OCPs were taking OCPs containing estrogen and progesterone (recorded/confirmed by staff during screening) >6 months and within 5-10 days of beginning a new pack. Sex workers not using hormonal contraceptives (NH) were not taking any hormonal contraception for >6 months and were within 5-10 days of beginning a menstrual cycle (proliferative phase). MPA was quantified in duplicate in the plasma of all women by ELISA (EuroProxima, Arnhem, Netherlands), as per protocol, following modifications (50) (Supplemental Figure 2A). Plasma concentrations of estradiol and progesterone were quantified for all women using the MILLIPLEX MAP Steroid/Thyroid Hormone Magnetic Bead Panel (Millipore, Merck KGaA, Darmstadt, Germany) and following the manufacturer’s instructions (Supplemental Figure 2B, C). Women were excluded if pregnant (or within 1 year), breastfeeding, unwilling to provide consent or follow protocol, using progesterone-only OCPs, or did not meet strict inclusion criteria.

At screening urine, blood, and vaginal swabs were collected. Urine was tested for *Neisseria gonorrhoeae*, and *Chlamydia* species by PCR (Xpert CT/NG kits, Cepheid AB, Solna, Sweden). Blood was collected for syphilis, and HIV serology, for all participants using a rapid test (Determine, Inverness Medical, Japan), and HIV serostatus was confirmed by ELISA (Vironostika, bioMérieux Clinical Diagnostics, Saint-Laurent, QC, Canada). Each woman underwent a gynaecological exam to obtain vaginal specimens for microscopy (Nugent Score, yeast infection, and *Trichomonas vaginalis*). Women positive for STI(s) were excluded and treated according to Kenyan protocols. HIV+ women were referred for anti-retroviral therapy.

Sample collection occurred within 1 week following screening. A PSA test (Seratec PSA Semiquant, Göttingen, Germany) was performed to confirm 12 hour abstinence, as recent unprotected sex could alter VMB. A gynecological exam was performed and endocervix washed with 2mL sterile 1X phosphate buffered saline (PBS) from a 3mL aliquot. CVL was collected from posterior vaginal fornix, placed in a sterile tube on ice, and sent to the laboratory where it was centrifuged to remove cellular debris. Supernatants and remaining 1mL PBS (negative controls) were aliquoted in a biosafety cabinet, frozen, and stored at −80°C until shipped in liquid nitrogen shipper to Winnipeg. CVLs and negative controls were shipped on dry ice to Hamilton for VMB analysis, α-amylase, and glycogen quantification. A cervical cytobrush to isolate cervical mononuclear cells (CMC) was collected following CVL and immediately sent to the University of Nairobi for flow cytometry.

### Bacterial V3 region of 16S rRNA gene sequencing

Bacterial DNA was extracted and purified from CVL supernatants as described (21). Four 1 mL aliquots of PBS (negative controls) were randomly selected for gDNA extraction and PCR amplification of the 16S rRNA gene. The hypervariable V3 region of 16S was amplified by PCR (21, 51). PCR products were sequenced by McMaster Genomics Facility (Hamilton, ON), Illumina MiSeq platform. Negative controls did not yield PCR products; bacterial contamination during sample collection, handling, processing, extraction, and PCR was considered negligible.

### Cytokine quantification

CVL cytokine concentrations were quantified by Milliplex MAP kit (Millipore, Billerica, MA) and analyzed by BioPlex-200 (BioRad, Mississauga, Canada). CVLs were incubated overnight, and analytes quantified included: IL-1α, IL-1ra, IL-1β, IL-8, IL-10, IL-17, sCD40L, IFN-γ, TNF-α, MIP-1α, MIP-1β, MIG-3, IP-10 and MCP-1.

### Glycogen and α-amylase quantification

Free glycogen in CVLs was quantified colorimetrically using the Glycogen Assay Kit (BioVision Inc., Milpitas, CA) (14). CVLs were thawed on ice, vortexed, and spun. CVL (5μl in quadruplicate) was added to a 96 well plate, and volume adjusted to 50μl with hydrolysis buffer. Hydrolysis enzyme was added to two wells/sample; those without enzyme were negative controls (background glucose in CVLs). Negative controls were subtracted from final values to determine total free glycogen in CVLs. ODs were read by SpectraMax i3 (Molecular Devices, Sunnyvale, USA). Alpha-amylase was quantified in CVLs (1:1 dilution, kit diluent) using a pancreatic human amylase ELISA (Abcam, Toronto, Canada) (52). Assay sensitivity is 4.0×10^-4^mg/mL (Biovision).

### Flow cytometry

CMC pellet was immunophenotyped at the University of Nairobi. Cells were washed (FACS Buffer) and stained for 30mins at 4°C with ECD-Live-Dead (Invitrogen, Carlsbad, USA) then washed twice with FACS. Cells were suspended in blocking solution (mouse IgG, FACS Buffer, FBS) for 10mins at 4°C and washed with FACS. A cocktail of PECy5-CD3, Alexa700-CD4, APC-H7-CD8, BB5151-CCR6, BV510-HLA-DR, PECy7-CD69, BV421-CCR5, APC-CD161, PE-CD38 and Brilliant Violet Stain (BD Biosciences, San Jose, USA) was used to stain cells for 30mins at 4°C. Cells were washed and fixed in 1% paraformaldehyde. Data were acquired on LSRII flow cytometer (BD System, San Jose, USA) and analyzed using FlowJo v10.0.8r1 (TreeStar, Ashland, USA).

### Humanized mice

Experimental protocols were approved by HiREB and McMaster University Animal Research Ethics Board (AREB), AUP# 14-09-40, in accordance with Canadian Council of Animal Care guidelines. Humanized NRG mice were generated as described (33). Briefly, 4 day-old mice were irradiated and injected with CD34-enriched placental cord blood stem cells, and left for 12 weeks to allow human immune reconstitution of bone marrow and peripheral tissues.

### Humanized mouse hormone treatments

Humanized mice (N=4) were anaesthetized and 21-day slow-release (476ng/mouse/day) estradiol pellets (Innovative Research of America, Sarasota, USA) were implanted in the neck (53). The quantity of estradiol induced is similar to the estrous cycle (54). To ensure subcutaneous injection of DMPA could induce circulating concentrations similar to women, we administered 1 (N=7) or 2mg (N=8) of DMPA or saline (control, N=5) to NRG mice, and collected blood by cardiac puncture 1 or 3 weeks later. MPA was quantified in duplicate in serum by ELISA (EuroProxima), as above (Supplemental Figure 2D). We used the 2mg dose in subsequent experiments because variability in circulating MPA was lower at 1 week, compared to 1mg dose, and because a similar peak and plateau phase occurred, as seen in women (4). Humanized mice (N=5) were administered 2mg DMPA (subcutaneous injection) (Pfizer, Mississauga, Canada). At baseline (day 0, prior to hormone treatment), 4, and 14 days under estradiol (N=4) or DMPA (N=5), vaginal bacteria were collected by anesthetizing (isoflurane), and washing vaginal tract 6X with 20μl sterile UV-treated PBS. Each 20μl was pipetted in/out of the vagina 8-10 times using sterile, filtered tips (BioRad). Care was taken to avoid contamination. Aliquots were frozen at −80°C until required.

### Bacterial sequencing of the humanized mouse VMB

Bacterial DNA was extracted and purified from 120μl of vaginal lavage following a modified protocol from Wessels et al., 2017 (21). Briefly, a nested PCR was performed using 5μl of template for 15 cycles using 8F and 1492R 16S primers. Then PCR product was used for a second reaction where 5μl of PCR product was run for 25 cycles using Illumina V3 primers (21). Samples were low biomass and negative ‘DNA extraction’ controls were carried out on reagents to ensure profiles were not due to contaminants.

### Glycogen quantification in the humanized mouse vagina

Free glycogen was quantified in vaginal tissue of humanized mice (N=10). Mice were treated with 2mg DMPA for 1 or 4 weeks (N=8, 4/time) as above and vaginal homogenates were boiled at 100°C for 10mins to inactivate enzymes, according to kit protocol. Homogenates (10μl in quadruplicate) were added to a 96 well plate, and assay performed as above.

### Intravaginal HIV-1 challenge in humanized mice

Control (N=20) and 2mg DMPA-treated (N=22) humanized mice were challenged intravaginally with 105 TCID50/mL NL4.3-Bal-Env HIV-1 (33). Humanized mice were HIV-1+ if viral load was detected in peripheral blood by clinical RT-PCR (33), at week 3 or 5 post-infection.

### Statistical analysis

16S rRNA sequences were processed by in-house data pipeline (21, 51). Alpha-diversity including singletons was calculated by sl1p pipeline (51), using QIIME version 1.7.0-dev. Ten rarefaction tables with 67821 sequences were used. Observed species, Chao1, and Shannon Diversity were graphed and analyzed using GraphPad Prism (GraphPad Software Inc., La Jolla, CA). Data are presented as mean±SEM. Linear discriminant analysis (LDA) effect size (LEfSe) (26) (https://huttenhower.sph.harvard.edu/galaxy/) was employed to determine significant taxonomic differences between groups. Alpha values of 0.05, and the 2.0 threshold for logarithmic LDA score for discriminative features were selected. Taxa bar charts, Bray-Curtis dissimilarity PCoAs, heatmaps, cluster analyses, cluster dendrograms, and species estimations were generated as described (21).

Categorical variables were compared by Fisher’s Exact Test or Chi-square (SigmaStat 3.5 Systat Software Inc., Chicago, IL, USA). Continuous variables were compared by student’s t-test, Mann-Whitney Rank Sum Test, one-way ANOVA, or Kruskal-Wallis one-way ANOVA (Graphpad Software Inc.). P≤0.05 was considered significant.

### Study approval

This study was approved by McMaster University (HiREB 0332-T) (AUP# 14-09-40), University of Manitoba (B2015:033), and Nairobi/Kenyatta National Hospital (KNH-ERC P132/03/2015). Written informed consent was received from all participants prior to inclusion in the study.

## Author Contributions

All authors contributed to interpretation of the data, drafting, and critically revising the manuscript. Conceived and Designed Study and Experiments: JMW, JL, MC, KRF, CK. Enrolled and Managed Study Participants: JL, KO, JK, JO, JC, JNM. Conducted Experiments: JMW, JL, MC, KO, AMF, DV, HAD, PVN, KM. Created the 16S Data Pipeline: MGS. Analyzed Data: JMW, MC, JL, KO, JCS, MGS. Created/provided Reagents and Humanized Mice: FV, AD, MJT, TM, AAA. Wrote the Manuscript: JMW, CK. Supervised the Study: JL, KO, JK, JO, JC, JNM, KRF, MGS, CK. **Competing Interests:** None to disclose.

## Supporting information

## Acknowledgments

The authors are sincerely grateful for all of the study participants, and would like to thank them for their participation in this trial. The authors would also like to acknowledge staff of the McMaster Central Animal Facility, and thank Laura Rossi, Michelle Shah, Dr. Fiona Whelan, and Ana Aquino for their technical assistance.

