## Supplementary material for "Medroxyprogesterone acetate mediated alteration in the vaginal microbiota and microenvironment in a Kenyan sex worker cohort"

### Supplemental Figure Legends:

#### Supplemental Figure 1: Experimental Design.

Flow chart depicting the experimental design, screening, and enrollment process for the Kenyan sex workers providing samples for the clinical study. CMC: cervical mononuclear cells. CVL: cervico-vaginal lavage. DMPA: Depot-medroxyprogesterone acetate. OCP: Oral contraceptive pills. PCR: polymerase chain reaction. PSA: prostate specific antigen.

#### Supplemental Figure 2: Circulating Hormones in Sex Workers and Humanized Mice.

**(A)** To ensure sex workers on DMPA had detectable amounts of MPA in the circulation, MPA was quantified in duplicate in the plasma of all study participants by ELISA. MPA was significantly higher in the sex workers in the DMPA group as compared to the OCP and no hormone groups (Kruskal-Wallis one-way ANOVA; $P \leq 0.0001$). **(B)** Additionally, sex workers on DMPA had significantly lower circulating estradiol than those not taking hormonal contraceptives (One-way ANOVA; $P \leq 0.05$). **(C)** Sex workers on DMPA also had significantly lower circulating progesterone than those on OCPs and not on hormonal contraceptives (Kruskal-Wallis one-way ANOVA; $P \leq 0.05$). **(D)** To determine if a subcutaneous injection of DMPA in the nape of the mouse neck could induce circulating concentrations similar to those observed in women, we administered 1 (N=7) or 2mg (N=8) of DMPA or saline as a control (N=5) to NRG mice (background strain of the humanized mice), collected peripheral blood by cardiac puncture 1 or 3 weeks later, and quantified MPA by ELISA. We proceeded to use the 2mg dose in subsequent humanized mouse experiments because there was less variability in circulating concentrations of MPA at 1 week as compared to the 1mg dose, and because this dose gave similar peak and plateau phase circulating concentrations to those observed in women (4). DMPA: Depot-medroxyprogesterone acetate. MPA: Medroxyprogesterone acetate. Mg: milligram. N/A: Not applicable. OCP: Oral contraceptive pills. ****: P≤0.0001. ***: P≤0.001. **: P≤0.01. *: P≤0.05. Data is presented as mean ± SEM.

**Supplemental Figure 3: DMPA Associated with Vaginal Bacterial Diversity in Sex Workers with Nugent Score ≤3.**

The Shannon Diversity Index was plotted in sex workers with Nugent Scores 0-3, to ensure the enhanced bacterial diversity we observed in the CVLs of sex workers on DMPA was not due to women with Nugent Scores 4-6. (**A**) Sex workers with Nugent Scores 0-3 who were on DMPA (N=17) had the greatest bacterial diversity followed by those with Nugent Scores 0-3 who were not on hormonal contraceptives (proliferative phase of the menstrual cycle, N=18), and those with Nugent Scores 0-3 on oral contraceptives (N=13), and at all levels of rarefaction in Shannon Diversity plots (P≤0.05; Kruskal-Wallis one-way ANOVA). (**B**) Sex workers with Nugent Scores 0-3 who were on DMPA had significantly greater bacterial diversity in their vaginal microbiota than those with Nugent Scores 0-3 who were not on hormonal contraceptives (proliferative phase of the menstrual cycle), at all depths of rarefaction (Mann-Whitney U test; P≤0.05). Even in this subset of sex workers with Nugent Scores of 0-3, the Shannon Diversity Index was significantly greater in sex workers on DMPA than those not on hormonal contraceptives. DMPA: Depot-medroxyprogesterone acetate. *: P≤0.05. **: P≤0.01. Data is presented as mean ± SEM.

**Supplemental Figure 4: Clustering of the Vaginal Microbiota in Sex Workers.**

**(A)** The gap statistic, which gives an estimation of how well ‘k’ (number of clusters) fits the data in the PCoA plot, was calculated. The number of clusters present in the data is indicated by the plateau in the gap statistic, which occurred at 3. **(B)** A heatmap of the top 20 species based on Bray-Curtis dissimilarity distance and PCoA ordination revealed that the vaginal microbiota (columns) did not cluster based on hormonal contraceptive type, but clustered by community state type (CST). Clustering of the vaginal microbiota by relative abundance of lactobacilli was also observed in the heatmap. Taxa are ordered along the y-axis by ranked order of abundance. CST: Community State Type. DMPA: Depot-medroxyprogesterone acetate. OCP: Oral contraceptive pills. *: resolved to bacterial genus.

**Supplemental Figure 5: Vaginal Glycogen and α-Amylase in Sex Workers on OCPs.**

As the diverse vaginal microflora can strongly influence vaginal carbohydrates in CVLs and overshadow the effect of hormones on these types of parameters (22, 23), we restricted our analysis of vaginal glycogen and α-amylase to sex workers with Nugent scores ≤3. **(A)** As estrogen is believed to enhance glycogen deposition in the vaginal epithelial cells, we quantified glycogen in the CVLs of sex workers on OCPs. Sex workers on OCPs had the highest levels of free vaginal glycogen, and significantly more free vaginal glycogen than the sex workers on DMPA (N=13, 16; P=0.008). **(B)** The quantity of α-amylase in the CVLs of sex workers on OCPs versus those on DMPA verged on significance (N=13, 16; P=0.11). DMPA: Depot-medroxyprogesterone acetate. OCP: Oral contraceptive pills.

**Supplemental Figure 6: Taxa Bar Charts of Humanized Mice Treated with E2 or DMPA over Time.**

The top 20 bacterial genera in the vaginal microbiota of humanized mice were plotted by relative abundance as taxa bar charts. Each bar represents the vaginal microbiota of one humanized mouse. Each colour represents a different genus of bacteria, as indicated in the legend. Each humanized mouse is denoted by a unique symbol above the taxa bar charts to allow for comparisons between days, and for comparisons with diversity plots in Figure 5 (E, F). Baseline vaginal washes were collected prior to hormone treatment while Days 4 and 14 were collected during hormone exposure. Although the bacterial species observed in the vaginal tract of humanized mice were different from those seen in women, hormones had a significant effect on the vaginal microbiota of humanized mice. **(A)** There was a significant reduction in diversity of the vaginal microbiota when humanized mice were treated with E2 (N=4), as demonstrated by taxa bar charts over time. Interestingly, administration of E2 rendered the typically diverse humanized mouse vaginal microbiota much more similar to the low-diversity, mono-colonized vaginal microbiota typically seen in women. **(B)** Diversity of the vaginal microbiota in humanized mice treated with DMPA (N=5) remained chronically high, suggesting that DMPA does not directly increase bacterial diversity within the vaginal microbiota, but rather that the presence of estradiol reduces bacterial diversity in the vaginal microbiota of humanized mice. E2: Estradiol. DMPA: Depot-medroxyprogesterone acetate. Hu-mouse: Humanized mouse. *: Resolved to family level; **: Resolved to order level.

### Supplemental Figures:

### Wessels et al., 2018 Supplemental Figure 1:

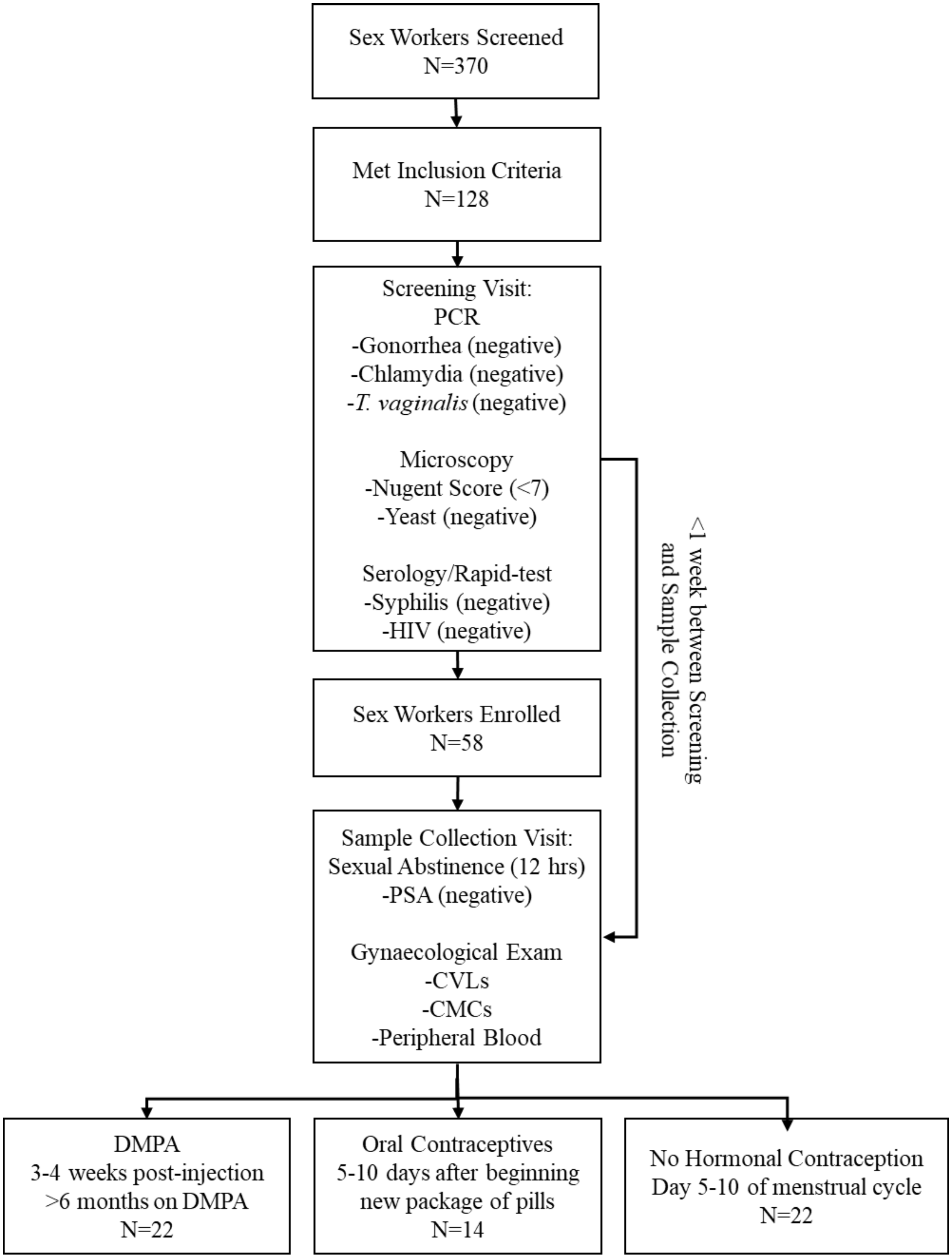

**Wessels et al., 2018 Supplemental Figure 2:**

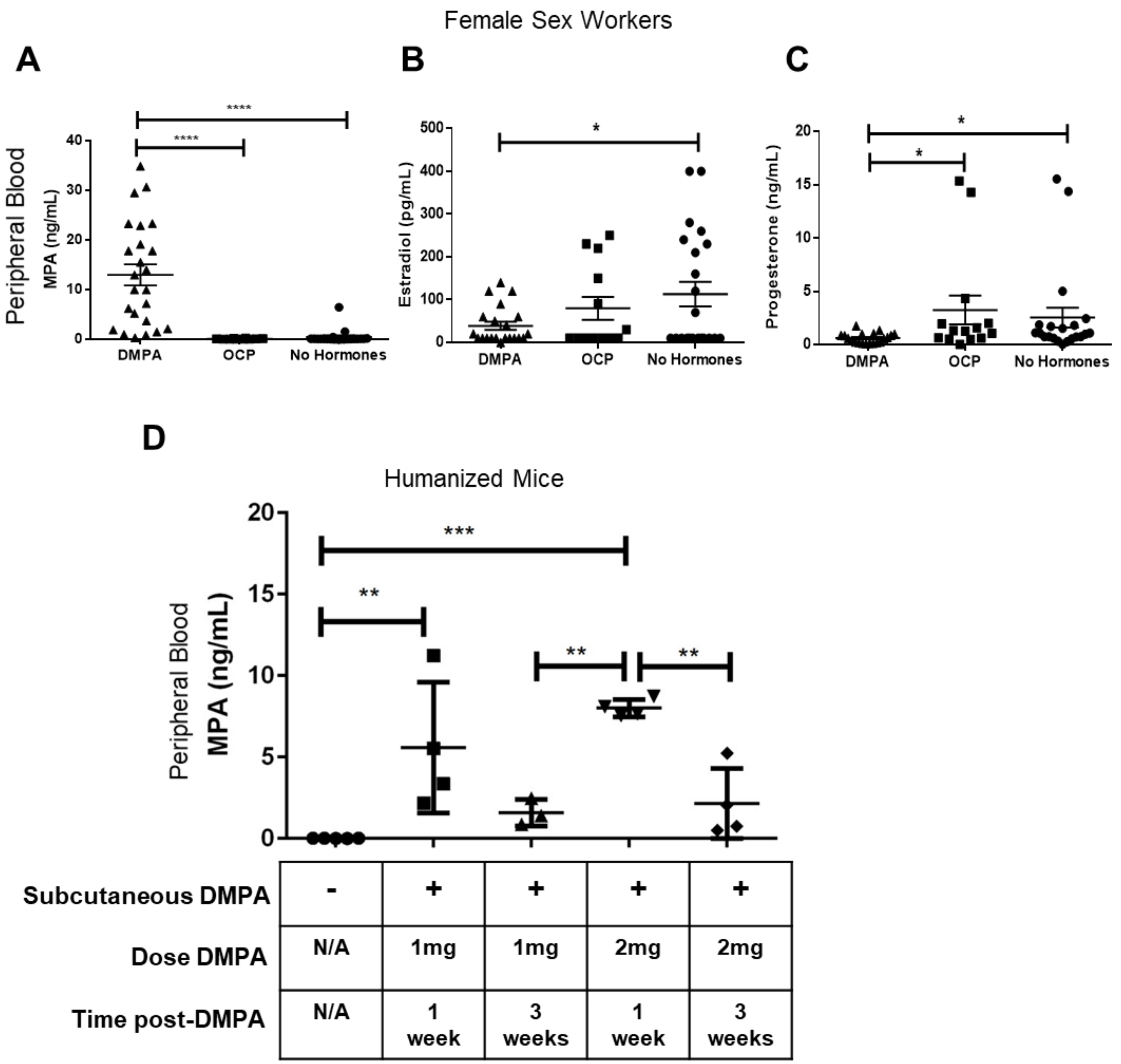

**Wessels et al., 2018 Supplemental Figure 3:**

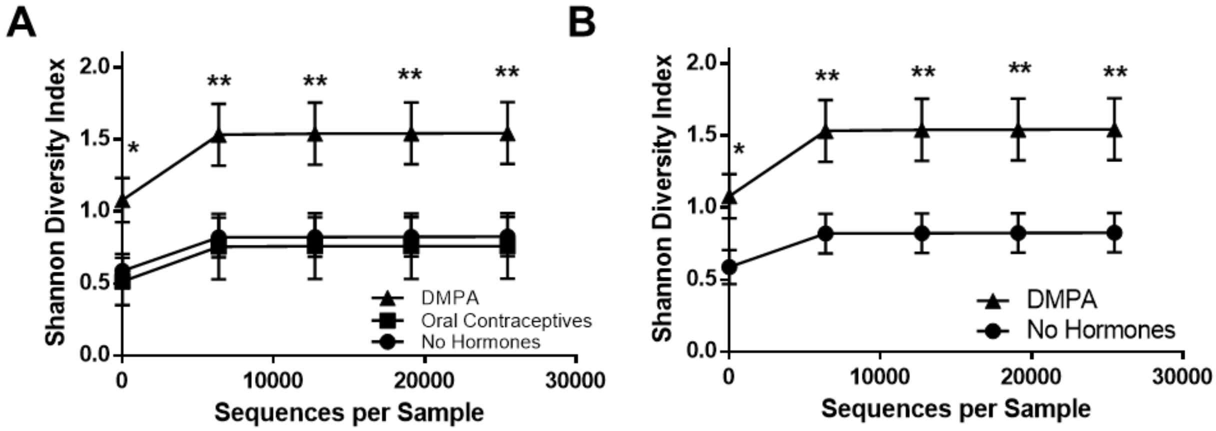

**Wessels et al., 2018 Supplemental Figure 4:**

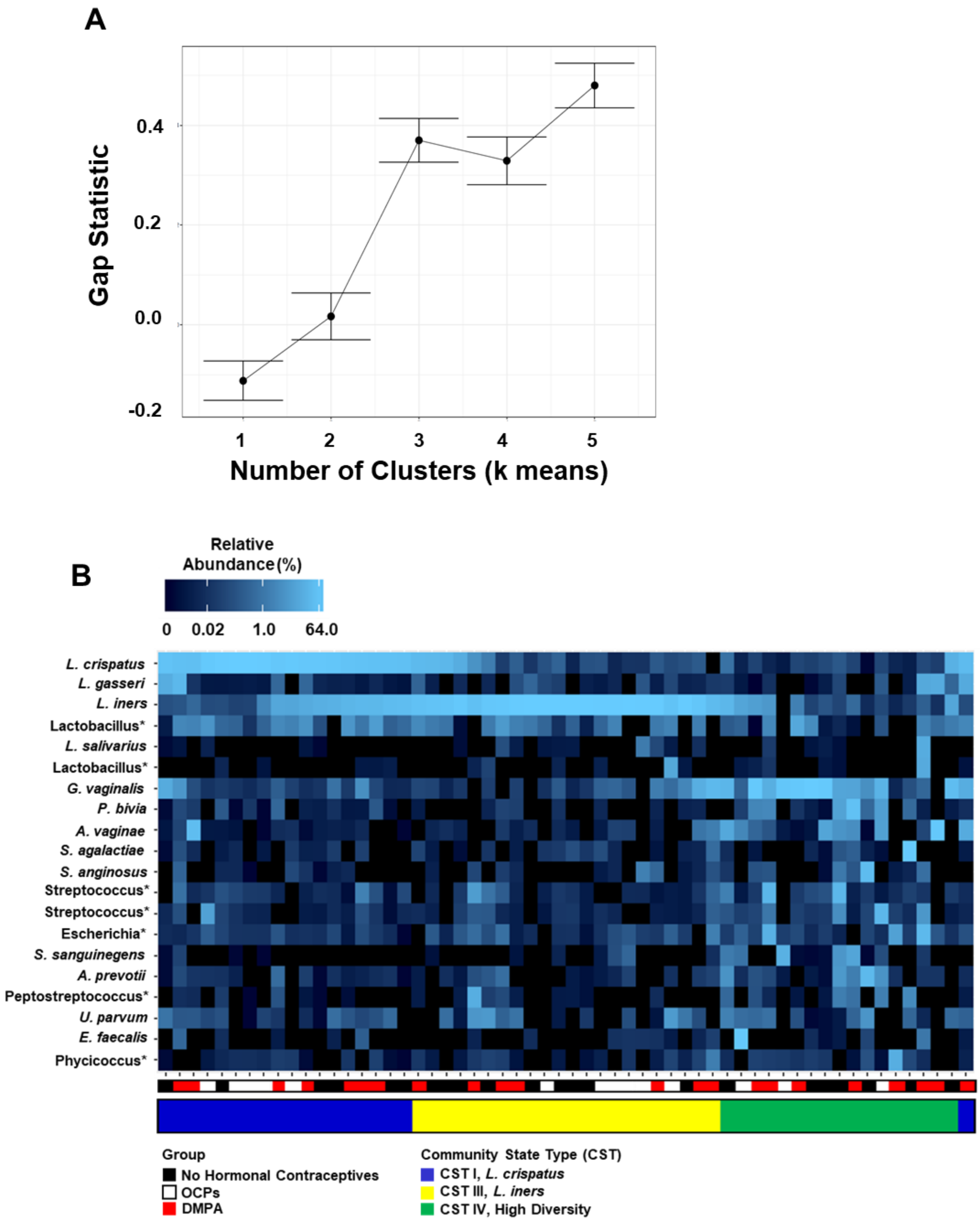

**Wessels et al., 2018 Supplemental Figure 5:**

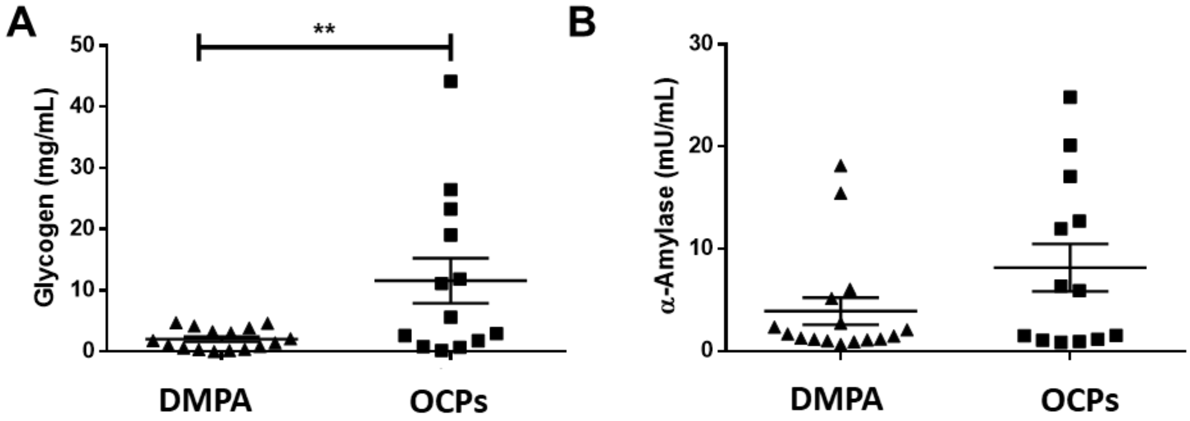

**Wessels et al., 2018 Supplemental Figure 6:**

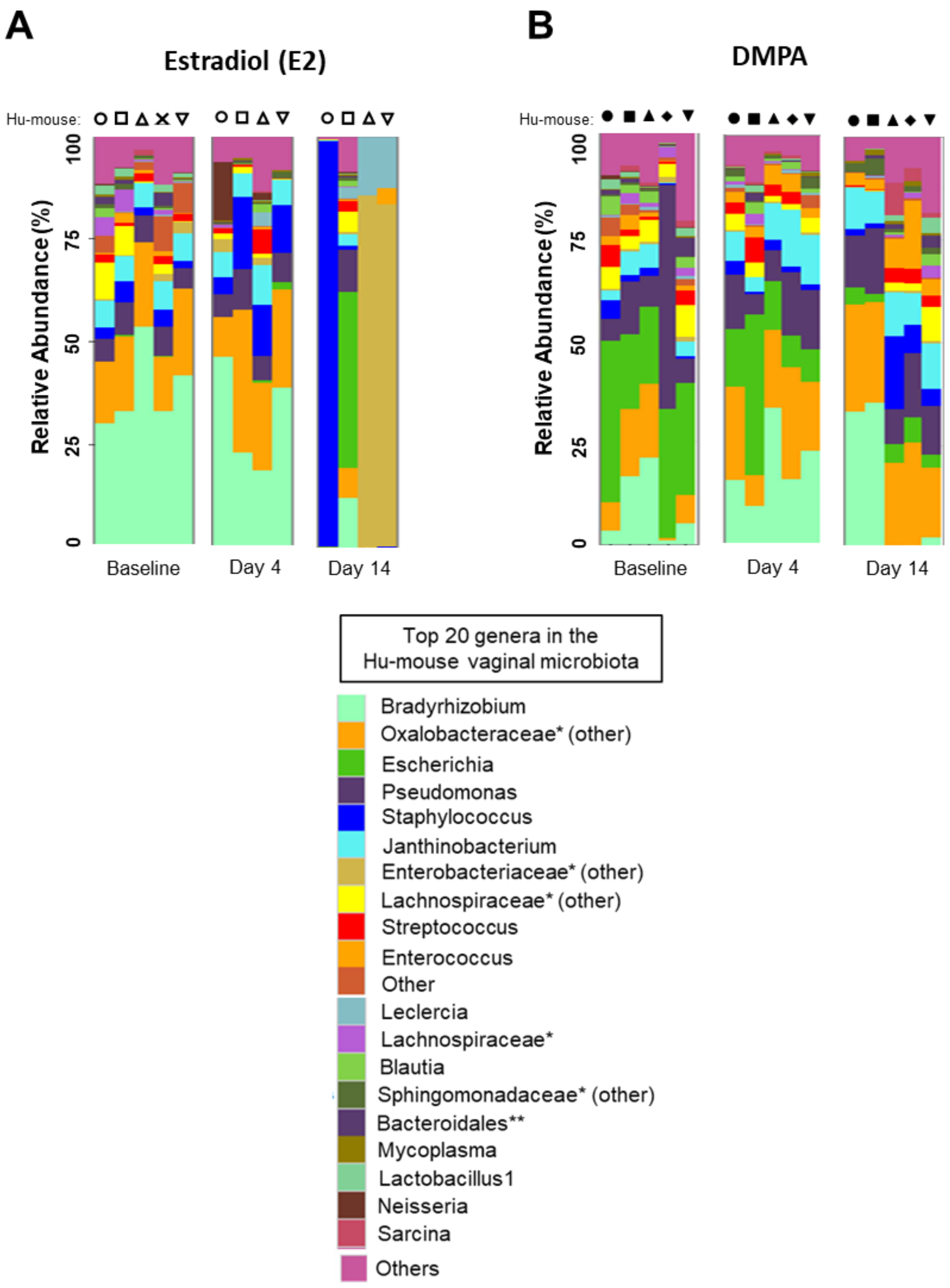
